# Default Mode and Salience Network Alterations in Suicidal and Non-Suicidal Self-Injurious Thoughts and Behaviors in Adolescents with Depression

**DOI:** 10.1101/2020.09.20.304204

**Authors:** Tiffany C. Ho, Johanna C. Walker, Giana I. Teresi, Artenisa Kulla, Jaclyn S. Kirshenbaum, Anthony J. Gifuni, Manpreet K. Singh, Ian H. Gotlib

## Abstract

Suicidal ideation (SI) and non-suicidal self-injury (NSSI) are two distinct yet often co-occurring risk factors for suicide in adolescents. Elucidating the neurobiological patterns that specifically characterize SI and NSSI in adolescents is needed to inform the use of these markers in intervention studies and to develop brain-based treatment targets. Here, we clinically assessed 70 adolescents—49 adolescents with depression and 21 healthy controls—to determine SI and NSSI history. Twenty-eight of the depressed adolescents had a history of SI and 29 had a history of NSSI (20 overlapping). All participants underwent a resting-state fMRI scan. We compared groups in network coherence of subdivisions of the central executive network (CEN), default mode network (DMN), and salience network (SN). We also examined group differences in between-network connectivity and explored brain-behavior correlations. Depressed adolescents with SI and with NSSI had lower coherence in the ventral DMN compared to those without SI or NSSI, respectively, and healthy controls (all *p*s<0.043). Depressed adolescents with NSSI had lower coherence in the anterior DMN and in insula-SN (all *p*s<0.030), and higher CEN–DMN connectivity compared to those without NSSI and healthy controls (all *ps*<0.030). Lower network coherence in all DMN subnetworks and insula-SN were associated with higher SI and NSSI (all *ps*<0.001). Thus, SI and NSSI are related to brain networks associated with difficulties in self-referential processing and future planning, while NSSI specifically is related to brain networks associated with disruptions in interoceptive awareness. Intrinsic network patterns may be reliable biomarkers of SI and NSSI in adolescents.

## Introduction

Suicide is currently the second leading cause of death in adolescents and young adults ages 15-24 years (1). While suicides are relatively rare, the prevalence of suicidal ideation (SI) and self-harming behaviors (including non-suicidal self-injury, or NSSI) among adolescents is alarmingly prevalent, with estimates as high as 38% (2, 3). SI and NSSI are also among the strongest clinical predictors of subsequent suicide attempt (4), which underscores the urgent need to advance our understanding of the neurobiological bases of these behaviors prior to an attempt in order to improve early detection and treatment.

Although SI and NSSI typically emerge during adolescence (2, 5) and often co-occur (6, 7), researchers and clinicians have long recognized that these phenomena are distinct (8, 9) and likely have unique neurobiological substrates (10, 11). Thus, elucidating the specific neurobiological patterns that characterize SI and NSSI in adolescents may shed insight into the diverse etiology of these behaviors, help to refine brain-based conceptual models (12, 11), and identify neural circuits that are sufficiently sensitive to assess the efficacy of targeted interventions (13).

Few researchers have used neuroimaging to examine SI or NSSI in adolescents, and almost no studies have utilized clinical controls (for reviews, 14, 11). To date, studies in this area have identified patterns of brain activation and connectivity related to SI and NSSI that are broadly consistent with neurobiological models of psychiatric disorders anchored in dysfunction of intrinsic large-scale brain networks (i.e., distributed regions with which intrinsic signals co-activate) of the human brain (15, 16): the central executive network (CEN), the default mode network (DMN), and the salience network (SN). The CEN is composed of dorsolateral prefrontal cortex (PFC) and posterior parietal regions that subserve working memory, executive attention, and other cognitive control functions. The DMN is composed of medial PFC, posterior cingulate cortex (PCC), precuneus, and hippocampus, and supports internally directed cognition and ruminative processing (17, 18). The SN is an integrative attentional system composed of the dorsal anterior cingulate cortex (dACC), anterior insula, amygdala, and striatum that evaluates and processes internal and external stimuli to facilitate the selection and deployment of appropriate behavioral responses (19, 20). While task-based fMRI studies have provided key knowledge of the psychological processes undergirding SI and NSSI, separately, in adolescents (11), these studies have not examined both SI and NSSI in a single cohort. Moreover, it is difficult to pinpoint the precise functional network patterns that are associated with these behaviors given the diverse tasks that have been used across investigations. Thus, it is imperative that we investigate patterns of intrinsic (i.e., task-independent) functional if we are to facilitate comparisons across samples and studies, and to identify specific neurobiological targets in clinical research studies.

In this context, advances in our understanding of intrinsic large-scale functional networks (21, 22) makes resting-state fMRI ideal for examining network-based perturbations in a variety of clinical conditions. Indeed, a small number of resting-state fMRI studies have documented nuanced alterations in the CEN, DMN, and SN in adolescents with history of suicide attempts, with SI, and with NSSI (for a review, see 11). In depressed adolescents, greater severity of SI has been associated with both greater connectivity between one node of the DMN (precuneus) with sensorimotor regions, as well as lower connectivity between another node of the DMN (PCC) with visual attention regions (23). In another sample of depressed adolescents, researchers documented lower network coherence (i.e., within-network connectivity) of the anterior node of the DMN (medial PFC), SN (centered on the dACC) and left CEN (24) in relation to participants’ most severe intensity of SI in the lifetime; further, longitudinal increases in network coherence of the SN (centered on the dACC) in this same sample were also associated with decreases in severity of SI (25). Compared to healthy controls, transdiagnostic samples of young adults (who were primarily diagnosed with depression) with history of SI were found to exhibit lower connectivity between the SN (dACC) and DMN (specifically, dorsal PCC relative to ventral PCC; 26). Finally, studies in adolescents and young adults with mood disorders with a history of attempt, who presumably have elevated SI, have found decreased and increased resting-state functional connectivity in different nodes of the DMN compared to non-attempters and healthy controls (27, 28, 29, 30, for a review, see 11).

Resting-state fMRI studies in adolescents with NSSI have also implicated many of the same networks. Compared to healthy controls, transdiagnostic samples of adolescents with NSSI had lower resting-state functional connectivity among nodes of the SN (amygdala, ACC, insula; 31), between regions of the SN and DMN (31), between the amygdala and supplemental motor area and visual attention regions (32), and between the ventral striatum and superior medial frontal cortex (33). Moreover, in studies examining treatments for NSSI in adolescents, improvement of NSSI was associated with lower amygdala-based resting-state functional connectivity with the SN (ACC), DMN (medial PFC), and higher amygdala-based resting-state functional connectivity with parahippocampal regions and temporal gyrus (31, 33).

Collectively, therefore, resting-state fMRI studies have evidenced primarily reduced within- and between-network connectivity of the CEN in relation to SI and NSSI, but with differential patterns of connectivity in the DMN and SN depending on the specific node being investigated. Although these networks are often treated as single functional entities, recent data-driven modeling approaches have demonstrated that these networks consist of distinct subnetworks with related but distinct functions (34, 35, 36). Moreover, all but two studies, which focused on depressed attempters (28, 29), did not include clinical controls (e.g., depressed adolescents without history of attempt). Study designs with clinical controls are necessary for the field to generate *specific*, in contrast to illness-general, biomarkers and to inform neurobiologically-based treatment targets for SI and NSSI in adolescents.

In the present study, we addressed these issues by examining differences in network coherence (i.e., within-network connectivity) of the CEN, DMN, and SN in depressed adolescents with suicidal ideation (SI+) compared to those without suicidal ideation (SI-) and psychiatrically healthy adolescents (CTL), as well as depressed adolescents with non-suicidal self-injury (NSSI+) compared to those without non-suicidal self-injury (NSSI-) and CTL. We conducted a comprehensive assessment of history of suicidal and self-harming thoughts and behaviors using several well-validated clinical interviews. In addition, we defined network coherence on the basis of resting-state fMRI timecourses, as these task-independent patterns of intrinsic functional signals are relatively stable and have strong test re-test reliability (37, 38, 39). Critically, we used a data-driven multivariate approach that is sensitive to detect the spatiotemporal subdivisions of the DMN and SN, circuits in which the greatest heterogeneity with respect to directionality and specific regions in resting-state and task-based fMRI findings have been reported (11). Based on the studies reviewed above, we hypothesized: 1) depressed adolescents with SI will exhibit lower CEN, DMN, and SN coherence compared to the other two groups and 2) depressed adolescents with NSSI will exhibit lower SN coherence compared to the other two groups. To complement these within-network connectivity analyses, we also examined group differences in between-network connectivity analyses. Finally, we also explored whether network coherence is associated with severity and frequency of SI and NSSI.

## Methods and Materials

### Participants and Study Design

Seventy-nine adolescents between the ages of 13 and 18 years were recruited from the San Francisco Bay Area community as part of a longitudinal study examining neurobiological mechanisms underlying adolescent stress and depression (K01MH117442). We interviewed participants at an initial session to assess study eligibility using the Kiddie Schedule for Affective Disorders and Schizophrenia–Present and Lifetime (K-SADS-PL; 40, 41), the Children’s Depressive Rating Scale–Revised (CDRS-R; 42), and the Family Interview for Genetics Studies (FIGS; 43). See “Clinical Assessments” in the Supplement for more details.

Inclusion criteria for potentially depressed adolescents included being between the ages of 13–18, fluency in English, and presence of a depressive disorder (Major Depressive Disorder or Dysthymia) based on a combination of the K-SADS-PL and CDRS-R. Inclusion criteria for potentially psychiatrically healthy adolescents included no current or lifetime diagnosis of any DSM-IV disorder or first-degree relative with history of suicide or known suicide attempt, or a probable diagnosis of depression, bipolar disorder, or schizophrenia using the FIGS. See “Inclusion/Exclusion Criteria” in the Supplement for more details.

Following the initial visit to determine participant eligibility through clinical interviews, participants were invited to return to the laboratory at a separate date to complete the MRI scan (interval between sessions: 11.07±6.02 days). Of the 79 adolescents enrolled in the study, 9 did not provide usable resting-state fMRI data for the present study (see *Resting-State fMRI Preprocessing*, below), resulting in a total of 70 adolescents for analyses: 49 depressed adolescents and 21 CTL. 28 depressed adolescents met criteria for clinically significant lifetime history of SI (SI+) and 21 did not (SI-) based on the K-SADS-PL. 29 depressed adolescents met criteria for clinically significant lifetime history of NSSI based on actions (NSSI+) and 20 depressed adolescents did not (NSSI-) based on the Self-Injurious Thoughts and Behaviors Interview (*SITBI*, see below). As expected, there was overlap in SI+ and NSSI+: 20 of the SI+ were also in the NSSI+ group. Because of the exploratory nature of the analyses conducted in this study, a formal power analysis was not conducted; however, previously published work in this area have had smaller sample sizes (23-26). The study was approved by the Institutional Review Board (IRB) at Stanford University. All participants and their parents gave written assent and informed consent, respectively, in accordance with the Declaration of Helsinki, and were financially compensated for their participation.

### C-SSRS

To comprehensively classify suicidal ideation and behavior, we administered the pediatric version of the Columbia Suicide Severity Rating Scale (C-SSRS; 44), which is a semi-structured interview that probes suicidal thoughts (including nature and severity of ideation) and behaviors (including preparatory acts, and actual, interrupted, or aborted attempts). Any actual, interrupted, or aborted attempt was classified as a suicide attempt.

### SITBI

To assess NSSI, we administered the Self-Injurious Thoughts and Behaviors Interview (SITBI; 45). The SITBI is a structured interview that assesses the history, frequency, and intensity of self-harming thoughts and behaviors that occurred without the intent to die, including lifetime frequencies and frequencies in the last year, month and week. To create a measure of current NSSI severity for correlational analyses, we used the sum of frequencies of thoughts and actions in the past month.

### SIQ-JR

To use a measure of current SI severity comparable to our measure of NSSI severity for correlational analyses, we administered the Suicidal Ideation Questionnaire–Junior (SIQ-JR; 46), a self-report 15-item measure rated on a 7-point scale, which evaluates frequency of SI in the past month.

### MRI Scanning Acquisition

All MRI scans were acquired at the Stanford Center for Cognitive and Neurobiological Imaging (CNI) with a 3T GE Discovery MR750 (General Electric Healthcare, Milwaukee, WI, USA) and Nova 32-channel head coil (Nova Medical, Wilmington, MA, USA). Participants completed a high-resolution T1-weighted MRI scan and an 8-minute T2*weighted resting-state fMRI scan with eyes closed. See “MRI Scanning Acquisition” in the Supplement for more details.

### Resting-State fMRI Preprocessing

As described in previous work across several independent samples (24, 25, 47, 48), structural and functional MRI data were preprocessed using an in-house script incorporating tools from FreeSurfer (49), FSL (50), and AFNI (51). For each subject, we computed mean relative displacement (MRD, also known as framewise displacement). Our criterion for excessive movement was defined as having more than 20 volumes with a MRD > 0.25 mm or a mean MRD > 0.2 mm (24, 25, 47, 48). No participants moved excessively based on these thresholds; however, 7 participants (2 CTL) did not complete the full scan due to discomfort and 2 participants (1 CTL) did not have physiological signal properly recorded. Participants who were included in our final analyses did not differ from these 9 participants on any demographic characteristics (all *p*s>0.259). See “Resting-State fMRI Preprocessing” in the Supplement for more details.

### Defining Functional Networks Using Group Independent Components Analysis (ICA)

We conducted a group-level independent component analysis (ICA) using FSL 6.0 MELODIC software version 3.14. Group ICA is a data-driven multivariate signal-processing method used to characterize spatiotemporal properties of functional MRI data (52, 53) that accounts for multiple voxel-voxel relations in order to define a spatial network of voxels based on the correlations of their timeseries. As in previous work (24, 25, 48), we identified a set of 25 probabilistic ICA components from the whole sample in order to compute individual-level metrics of network coherence from a comparable spatial set of components for further statistical analysis (see *Network Coherence*, below, for more details). Based on our hypotheses, we focused on the subdivisions of the CEN, SN, and DMN. Seven networks of interest were identified on the basis of their neuroanatomical components by trained raters (TCH, JSK, AG): left and right CEN (CEN-L, CEN-R), centered on frontoparietal regions, anterior subdivision of the DMN centered on mPFC (DMN-A), posterior subdivision of the DMN centered on the PCC (DMN-P), ventral subdivision of the DMN centered on the middle temporal lobe, hippocampus, and parahippocampal region (DMN-V), a subdivision of the SN centered on the dorsal anterior cingulate cortex and anterior insula (SN-ACC), and a subdivision of the SN centered on the mid- and posterior insula, striatum, and somatosensory cortex (SN-Ins). See Figure 1 for more details.

**Figure 1.**
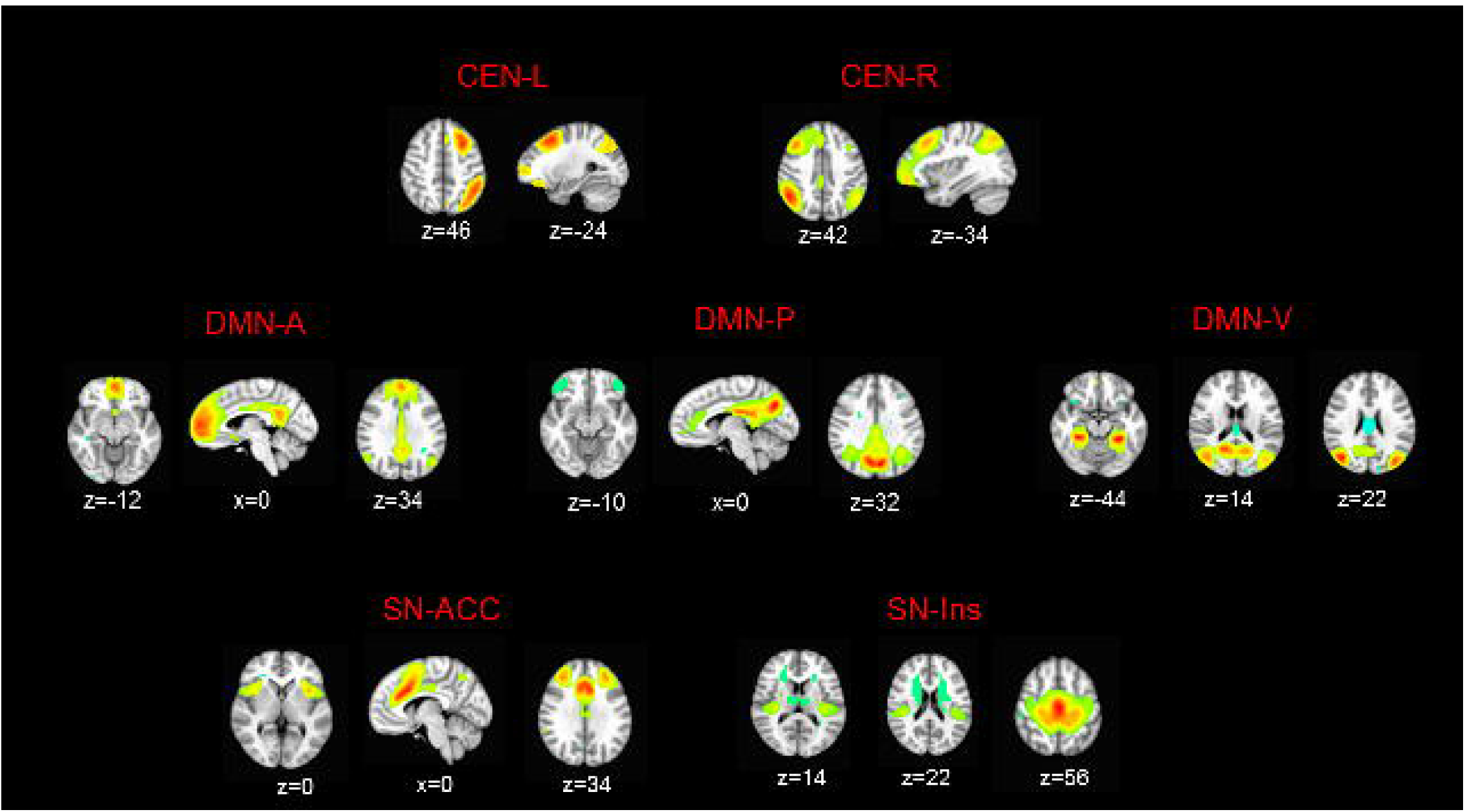
Intrinsic networks derived from group ICA. Maps were thresholded at *t*(68)>3.93 (α = 0.0001). All views and coordinates are presented in radiological convention. CEN includes dorsolateral prefrontal cortex, middle frontal gyrus, and inferior parietal lobule, DMN-A includes anterior medial prefrontal cortex, subgenual anterior cingulate cortex, and cingulate gyrus, DMN-P includes posterior cingulate cortex, precuneus, and inferior parietal lobule, DMN-V includes middle temporal lobe, hippocampus/parahippocampal region, and posterior cingulate cortex, SN-ACC includes dorsal anterior cingulate cortex and anterior insula, SN-Ins includes mid- and posterior insula, striatum, and somatosensory cortex; CEN-L=left central executive network; CEN-R=right central executive network; DMN-A=anterior default mode network; DMN-P=posterior default mode network; DMN-V=ventral default mode network; SN-ACC=dorsal anterior cingulate cortex-based salience network; SN-Ins=insula-based salience network.

### Network Coherence

We operationalized network coherence as the strength of the associations among timecourses of all voxels within a given network for each individual (24, 25, 48). Specifically, we applied dual regression to generate individual-level spatial maps by regressing group-level spatial maps onto each individual’s 4D dataset, resulting in a set of individual-level timeseries data per spatial component. Each individual timeseries was then regressed on the same 4D dataset, resulting in a set of individual-level spatial components, wherein each voxel contains a regression weight reflecting the strength of functional connectivity of its timeseries with the identified network while controlling for associations from all other networks. These timeseries, within each voxel, were then z-scored (i.e., normalized by the residual within-subject variance). We then averaged the voxel-wise z-scores within each network component (defined from the group-level maps to facilitate comparability) and used this value as an index of network coherence.

### Between-Network Connectivity

To complement our within-network analyses, we performed seed-to-seed connectivity using the networks defined by the ICA analysis and computed the mean temporal correlation across all voxels within each network. Based on our results (see *Results*, below), we focused on connectivity between CEN and SN-ACC, as well as each of these networks with the three DMN networks.

### Statistical Analyses

All statistical analyses were conducted in R (v 3.5.3; 54) and all significance tests were two-tailed. Levene’s test was used to determine homogeneity of variance; non-parametric tests were used in all instances where groups differed in homogeneity of variance and also for NSSI frequency of thoughts and actions because of the skewness of those data (see Figure S1). Because CTL participants were included in testing differences in diagnostic groups, we did not include depression severity scores due to statistical collinearity. Given evidence of sex differences and age-related effects in both SI and NSSI behaviors (2,5), we included age and sex (dummy coded as a binary factor), as well as relevant covariates that may affect patterns of resting-state functional connectivity (i.e., motion during the resting-state fMRI scan as measured by mean relative displacement, and psychotropic medication use, dummy coded as a binary factor). All covariates were mean-centered. Following the detection of a significant group effect, we conducted planned linear contrast tests to examine differences in SI+ and NSSI+ from their respective clinical and healthy counterparts in the following networks: CEN-L, CEN-R, DMN-A, DMN-P, DMN-V, SN-ACC, and SN-Ins. Because of our planned contrasts, we determined significance at *p*<0.05, uncorrected; however, we also report FDR-corrected significance values for the group effect (SI and NSSI comparisons, separately). Because we did not find group differences in CEN and SN-ACC within-network coherence (see *Results*, below), we also conducted linear regressions to test for group differences in between-network connectivity among these networks and with each of these networks with the three DMN networks. Among the depressed adolescents only, we explored associations in network coherence with severity of SI in the past month, and with frequency of NSSI thoughts and actions in the past month.

### Code Availability

Scripts for preprocessing the resting-state fMRI data and for conducting statistical analyses can be found at: https://github.com/tiffanycheingho/TIGER/.

## Results

### Demographic and Clinical Characteristics

Demographic and clinical characteristics for the depressed and CTL, SI+ and SI-, and NSSI+ and NSSI-participants are presented in Tables 1ABC, respectively. As expected, the depressed and CTL adolescents differed significantly in clinical characteristics including CDRS-R scores, SIQ scores, and medication status (all *ps<*0.0002). The depressed and CTL adolescents did not differ in any demographic characteristics (all *p*s>0.363) except for motion (*p*=0.040), where CTL adolescents moved more, on average. See Table 1A for more details.

**Table 1.**
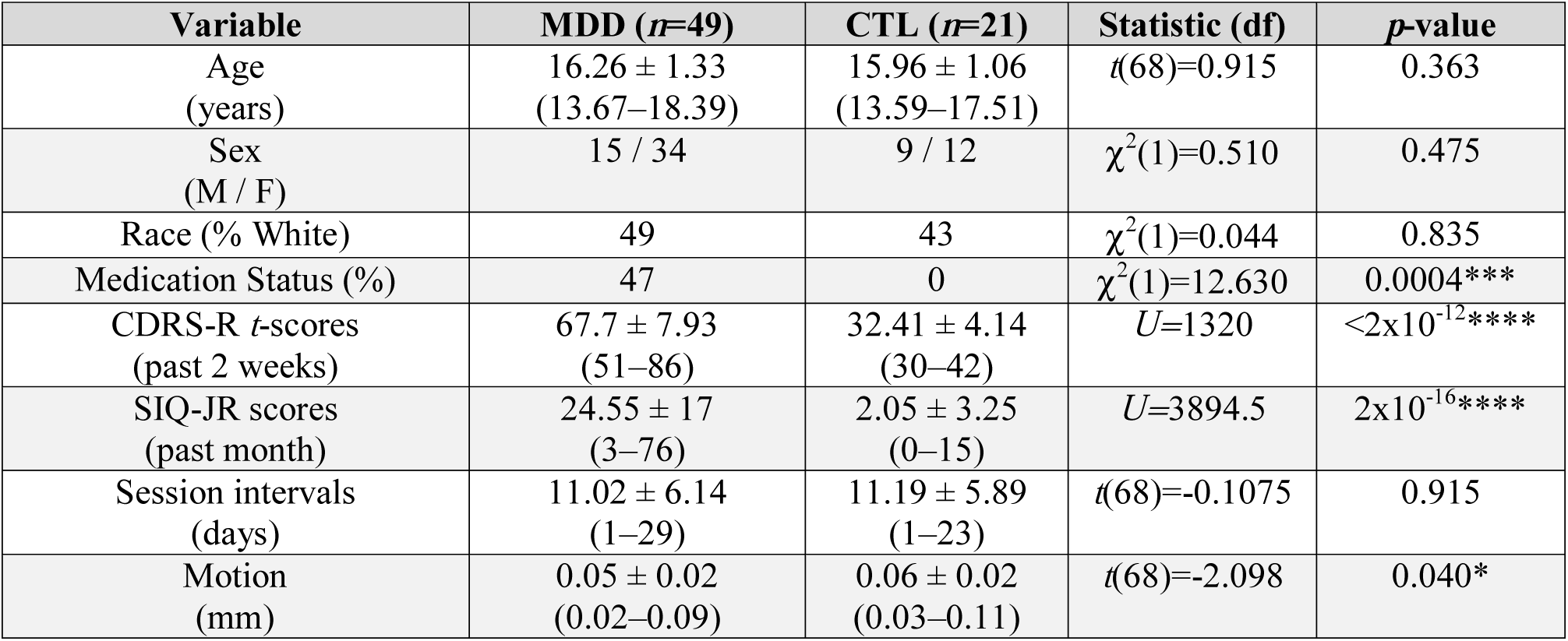
Descriptive statistics for depressed and non-depressed adolescents. Continuous variables are reported as mean ± SD (min – max). Numbers in [] indicate number of missing participants. See Tables 1B and 1C for descriptive statistics for SI+ versus SI- and NSSI+ versus NSSI-, respectively. CDRS-R=Children’s Depressive Rating Scale-Revised; MDD=Major Depressive Disorder or Dysthymia; SIQ-JR=Suicidal Ideation Questionnaire-Junior. \**p*<0.05, \*\**p*<0.01, \*\*\**p*<0.001, \*\*\*\**p*<0.0001

**Table 1.**
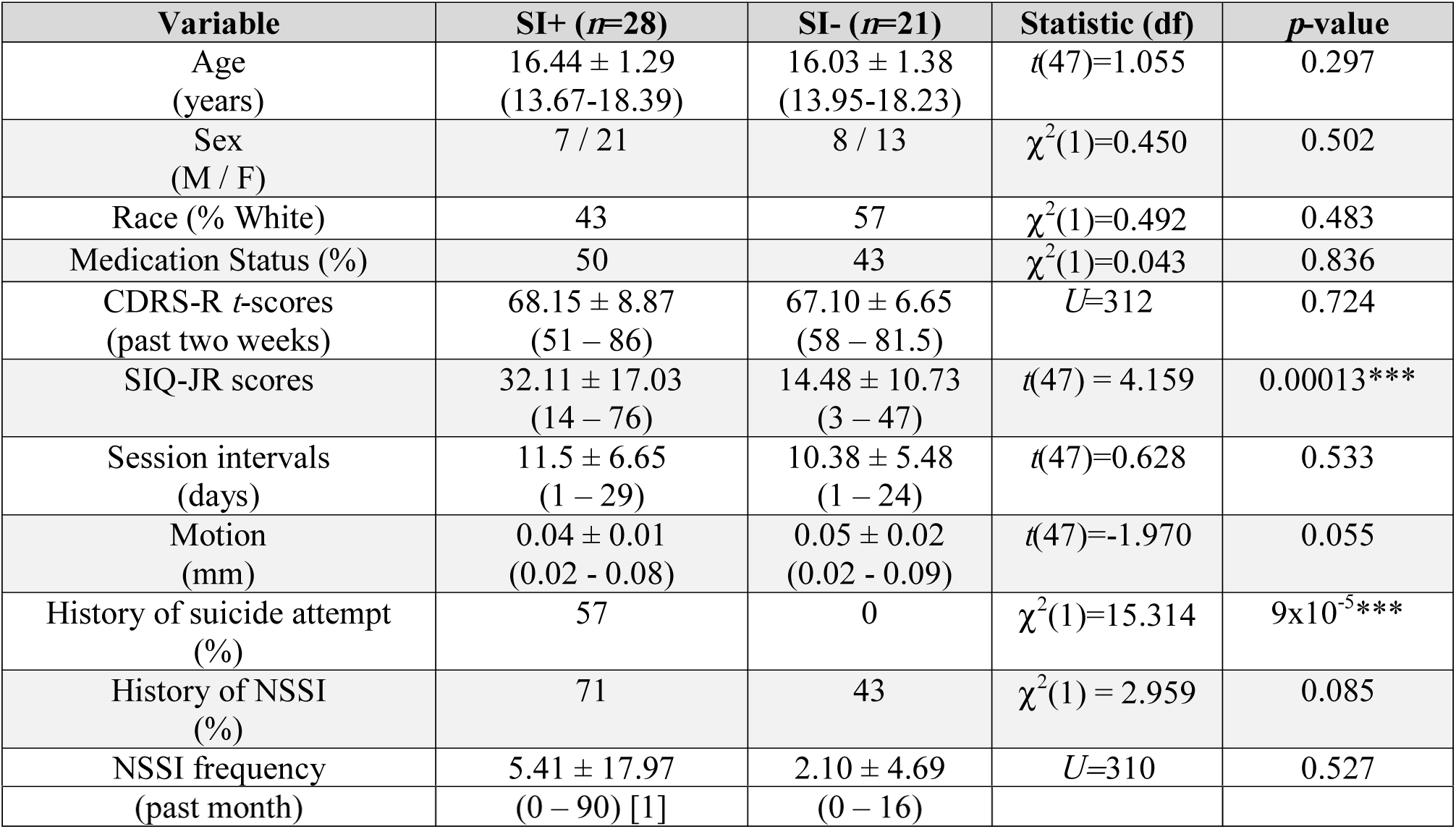
Descriptive statistics for depressed with suicidal ideation (SI+) and without suicidal ideation (SI-). Continuous variables are reported as mean ± SD (min – max). Numbers in [] indicate number of missing participants. CDRS-R=Children’s Depressive Rating Scale-Revised; SIQ-JR=Suicidal Ideation Questionnaire-Junior. \**p*<0.05, \*\**p*<0.01, \*\*\**p*<0.001, \*\*\*\**p*<0.0001

**Table 1.**
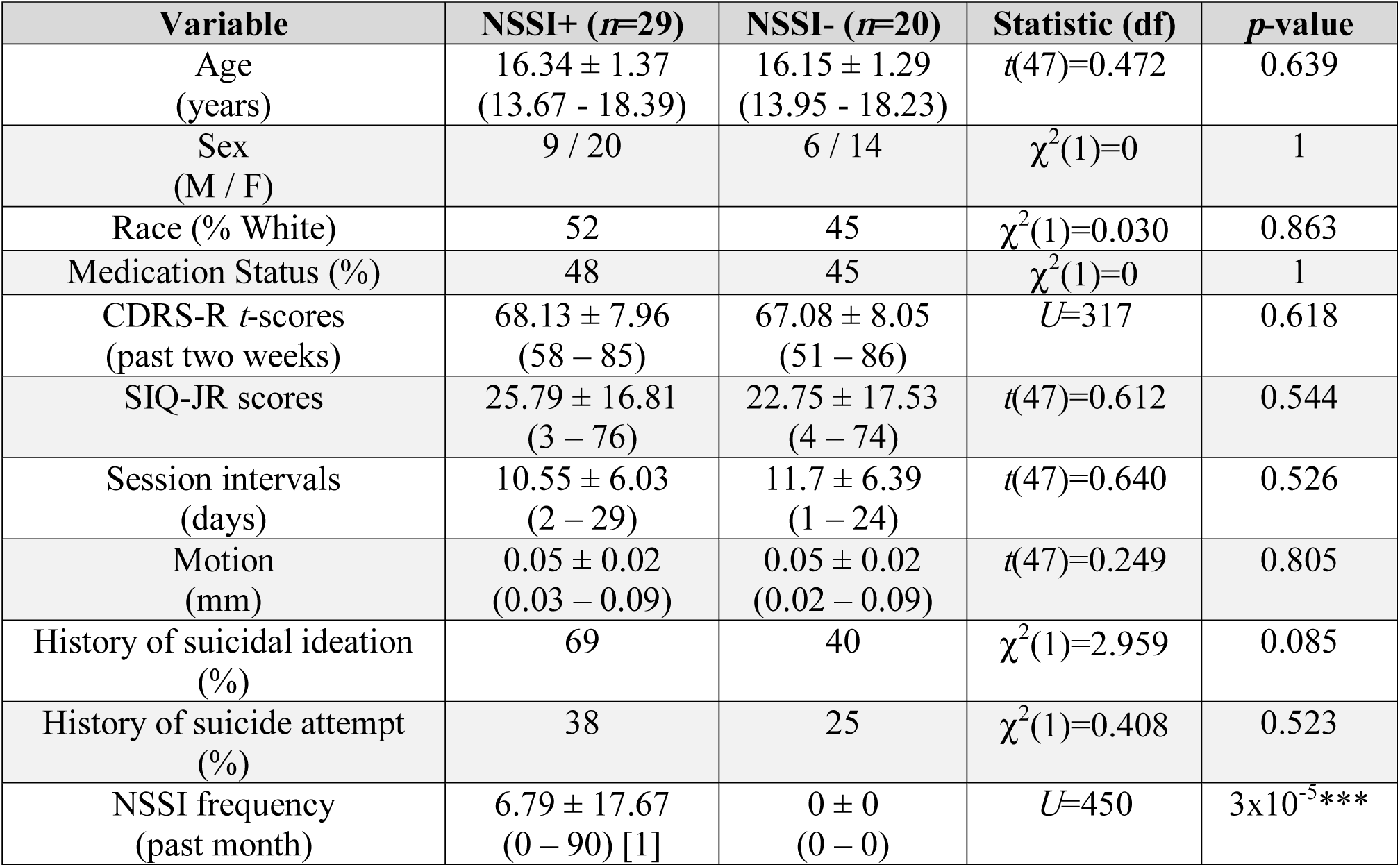
Descriptive statistics for depressed with non-suicidal self-injury (NSSI+) and without non-suicidal self-injury (NSSI-). CDRS-R=Children’s Depressive Rating Scale-Revised; SIQ-JR=Suicidal Ideation Questionnaire-Junior. \**p*<0.05, \*\**p*<0.01, \*\*\**p*<0.001,

As expected, SI+ and SI-participants differed significantly in history of suicide attempt (57% of SI+ had a history of attempt and 0% in SI-) and SIQ scores (all *p*s<0.00013). Critically, SI+ and SI-participants did not differ in medication status, CDRS-R scores, and in history or frequency of NSSI (all *p*s>0.085). See Table 1B for more details. Finally, as expected, NSSI+ and NSSI-participants differed significantly in frequency of NSSI thoughts and actions in the past month (*U*=450; *p*=3.23×10^−5^). Critically, NSSI+ and NSSI-did not differ in medication status, CDRS-R scores, history of suicide attempt or SI, or SIQ scores (all *p*s>0.085). See Table 1C for more details.

### Network Coherence Differences Associated with SI

To investigate patterns of network coherence associated with SI, we conducted linear models predicting network coherence values based on group (SI+, SI-, CTL) while adjusting for covariates. SI+ had significantly lower coherence in the DMN-V compared to both SI-(*p*=0.043) and CTL (*p*=0.021); SI-did not differ from CTL in coherence of the DMN-V (*p*=0.601). While SI+ and SI-participants did not differ in coherence of DMN-A, DMN-P, and SN-Ins (all *p*s>0.280), SI-had significantly lower coherence in DMN-P (*p*=0.038) compared to CTL. See Table 2 and Figure 2 for more details.

**Table 2.**
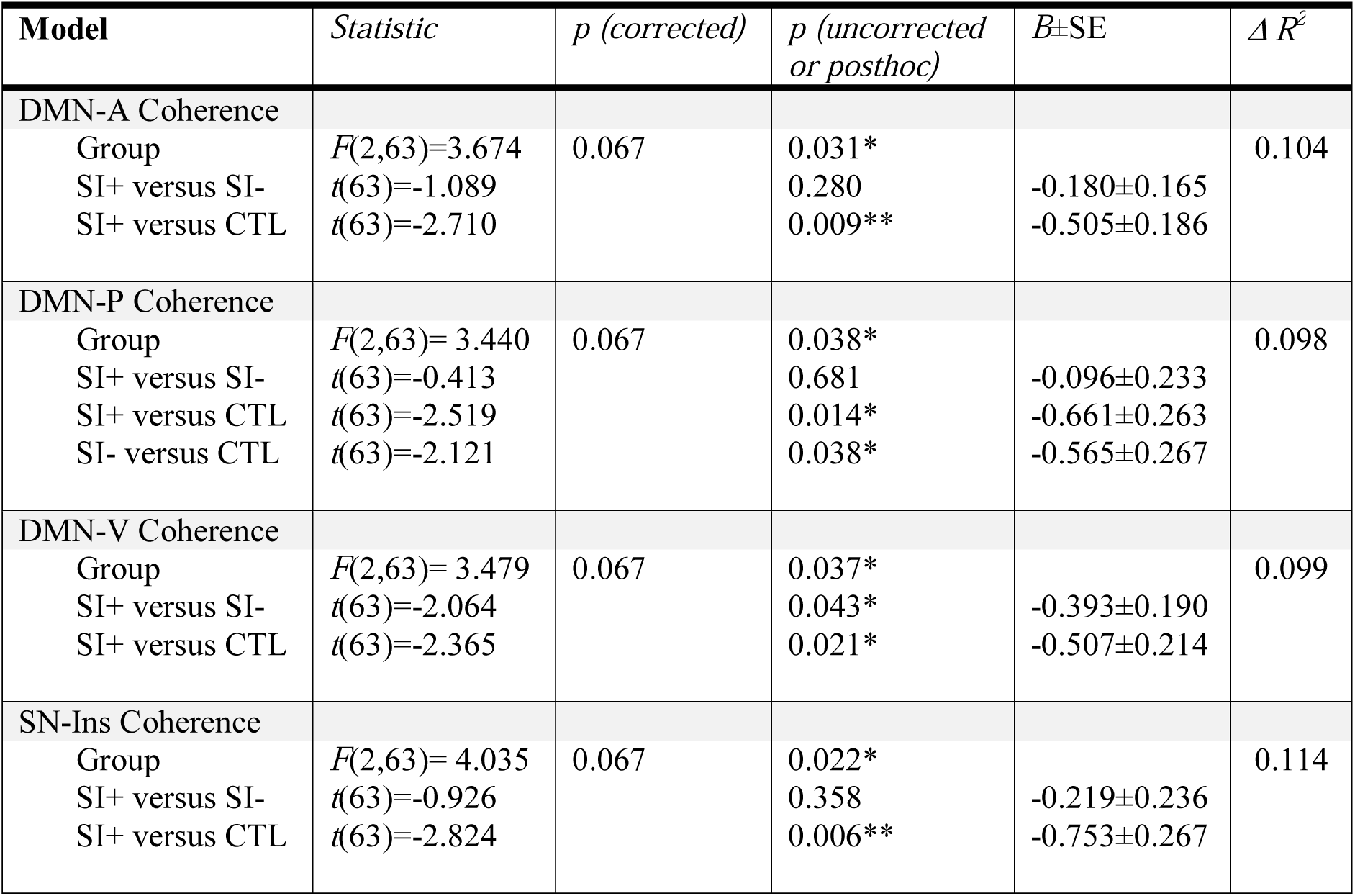
Results of statistical models with a significant overall effect of group (SI+, SI-, CTL) on network coherence with planned linear contrast tests (SI+ versus SI- and SI+ versus CTL). Posthoc linear contrast tests (SI-versus CTL) with significant results are also reported. All results shown include age, sex, motion, and medication use as covariates (see Table S1 for more details). See Figure 2 for more details. FDR-corrected *p*-values are reported for the main effect of group only. \**p*<0.05, \*\**p*<0.01. SI=suicidal ideation; CTL=healthy control

**Figure 2.**
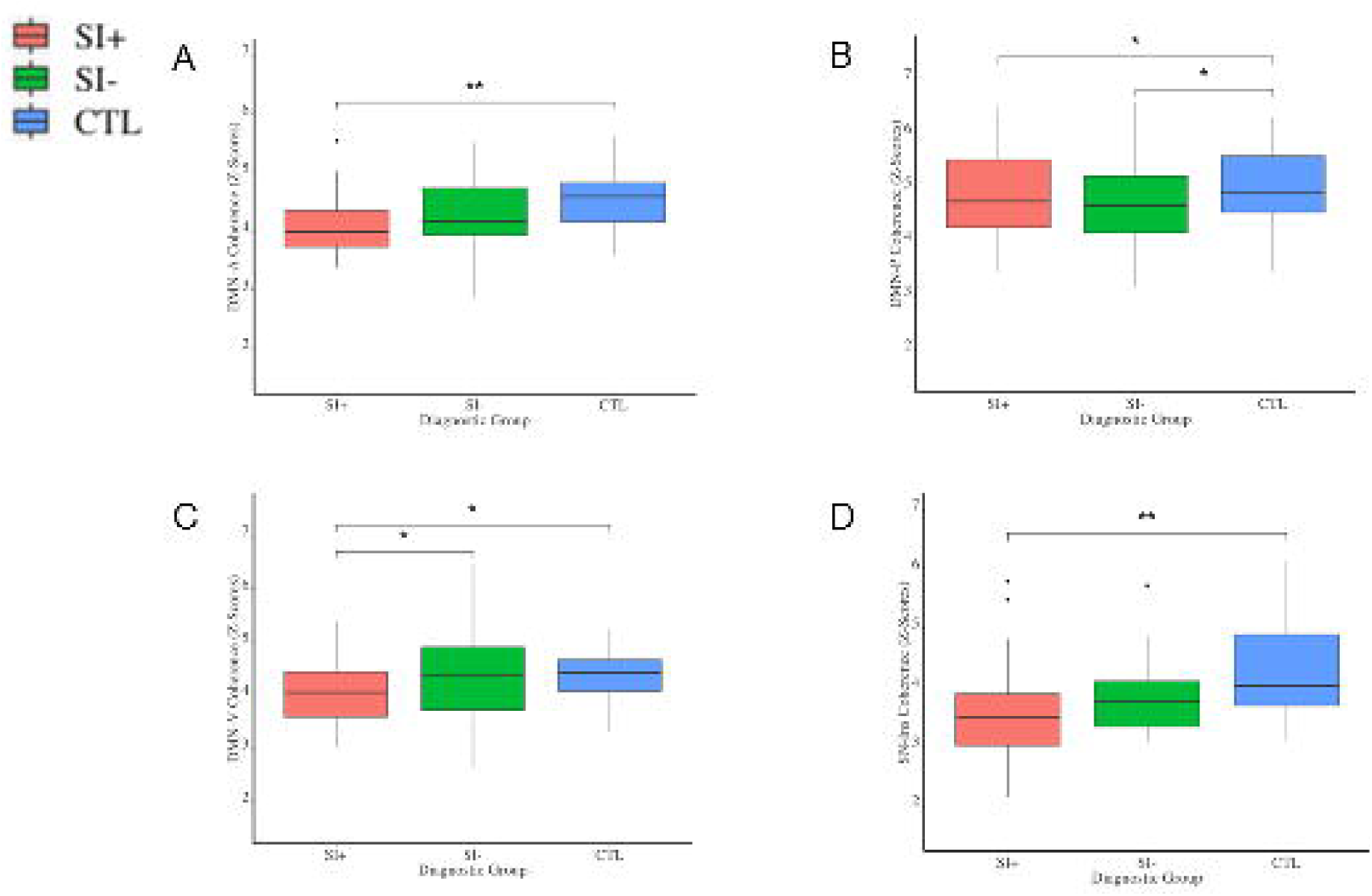
Network coherence patterns associated with suicidal ideation. SI+ exhibited significantly lower network coherence in DMN-A **(A)**, DMN-P **(B)**, DMN-V **(C)**, SN-Ins **(D)** compared to CTL. SI+ exhibited significantly lower network coherence in DMN-V compared to SI-(**C**). SI-exhibited significantly lower network coherence in DMN-P compared to CTL (**B**). For the purposes of visualization, all box plots depict raw data without adjustment of covariates. See Tables 2 and S1 for more details. DMN-A=anterior default mode network; DMN-P=posterior default mode network; DMN-V=ventral default mode network; SI=suicidal ideation; CTL=healthy controls. \**p*<0.05, \*\**p*<0.01 (uncorrected)

### Network Coherence Differences Associated with NSSI

To investigate patterns of network coherence associated with NSSI, we conducted linear models predicting network coherence from group (NSSI+, NSSI-, CTL) while adjusting for covariates. NSSI+ had significantly lower network coherence in the DMN-A (*p*=0.030), DMN-V (*p*=0.017), and SN-Ins (*p*=0.001) compared to NSSI-, and significantly lower network coherence in the DMN-A (*p*=0.0007), DMN-V (*p*=0.026), and SN-Ins (*p*=0.0006) compared to CTL; NSSI-did not differ significantly from CTL in coherence of these networks (all *p*s>0.147). NSSI+ participants had significantly lower network coherence in DMN-P compared to CTL (*p*=0.002) but did not differ from NSSI-(*p*=0.297); NSSI-also exhibited significantly lower network coherence in the DMN-P compared to CTL (*p*=0.034). See Table 3 and Figure 3 for more details. SI+ and NSSI+ did not differ significantly in network coherence in CEN-L, CEN-R, and SN-ACC from their clinical counterparts or healthy controls (all *p*s>0.155).

**Table 3.** Results of statistical models with a significant overall effect of group (NSSI+, NSSI-, CTL) on network coherence with planned linear contrast tests (NSSI+ versus NSSI- and NSSI+ versus CTL). Posthoc linear contrast tests (NSSI-versus CTL) with significant results are also reported. All results shown include age, sex, motion, and medication use as covariates (see Table S2 for more details). See Figure 3 for more details. FDR-corrected *p*-values are reported for the main effect of group only. \**p*<0.05, \*\**p*<0.01, \*\*\**p*<0.001. NSSI=non-suicidal self-injury; CTL=healthy control

| Model | Statistic | $p$ (corrected) | $p$ (uncorrected or posthoc) | $B \pm SE$ | $\Delta R^2$ |
| --- | --- | --- | --- | --- | --- |
| DMN-A Coherence |  |  |  |  |  |
| Group | $F(2,63)=6.835$ | 0.007 | 0.002** | | 0.178 |
| NSSI+ versus NSSI- | $t(63)=-2.223$ | | 0.030* | -0.348±0.157 | |
| NSSI+ versus CTL | $t(63)=-3.568$ | | 0.0007*** | -0.622±0.174 | |
| DMN-P Coherence |  |  |  |  |  |
| Group | $F(2,63)=5.333$ | 0.016 | 0.007** | | 0.145 |
| NSSI+ versus NSSI- | $t(63)=-1.052$ | | 0.297 | -0.236±0.225 | |
| NSSI+ versus CTL | $t(63)=-3.261$ | | 0.002** | -0.814±0.250 | |
| NSSI- versus CTL | $t(63)=-2.165$ | | 0.034* | -0.578±0.267 | |
| DMN-V Coherence |  |  |  |  |  |
| Group | $F(2,63)=4.102$ | 0.037 | 0.021* | | 0.115 |
| NSSI+ versus NSSI- | $t(63)=-2.456$ | | 0.017* | -0.459±0.187 | |
| NSSI+ versus CTL | $t(63)=-2.279$ | | 0.026* | -0.473±0.207 | |
| SN-Ins Coherence |  |  |  |  |  |
| Group | $F(2,63)=8.833$ | 0.003 | 0.0004*** | | 0.219 |
| NSSI+ versus NSSI- | $t(63)=-3.329$ | | 0.001** | -0.731±0.220 | |
| NSSI+ versus CTL | $t(63)=-3.618$ | | 0.0006*** | -0.883±0.244 | |

**Figure 3.**
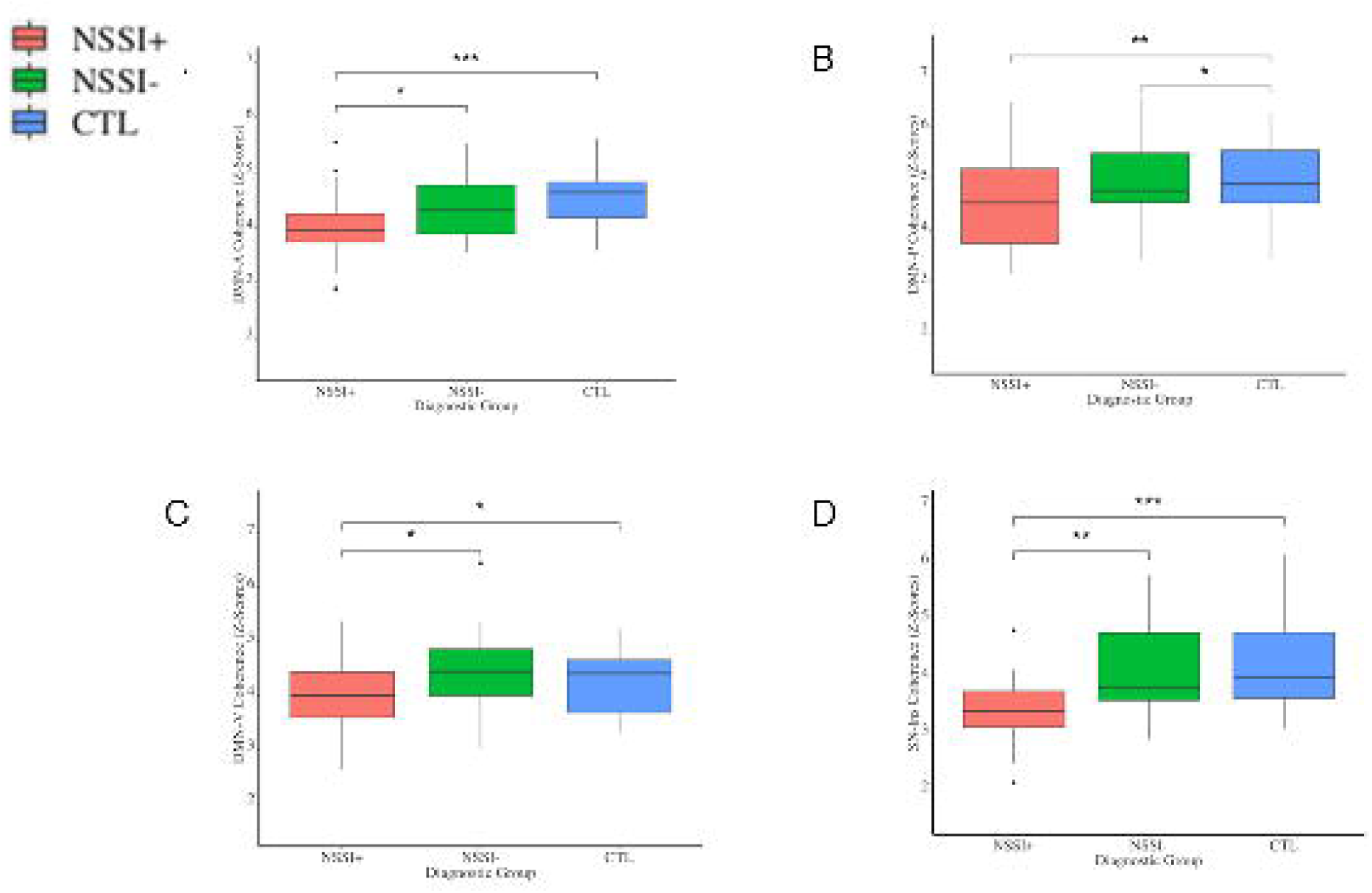
Network coherence patterns associated with non-suicidal self-injury. NSSI+ exhibited significantly lower network coherence in DMN-A **(A)**, DMN-P **(B)**, DMN-V **(C)**, SN-Ins **(D)** compared to CTL. NSSI+ exhibited significantly lower network coherence in DMN-A **(A)**, DMN-V **(C)**, and SN-Ins **(D)** compared to NSSI-. NSSI-exhibited significantly lower network coherence in DMN-P (**B**) compared to CTL. For the purposes of visualization, all box plots depict raw data without adjustment of covariates. See Tables 3 and S2 for more details. DMN-A=anterior default mode network; DMN-P=posterior default mode network; DMN-V=ventral default mode network; NSSI=non-suicidal self-injury; CTL=healthy controls. \**p*<0.05, \*\**p*<0.01, \*\*\**p*<0.001 (uncorrected)

### Between-Network Connectivity Results

To complement our primary investigation examining network coherence (i.e., within-network connectivity), we tested whether groups differed in between-network connectivity of the CEN and SN-ACC with the three subdivisions of the DMN, which were the networks we did not find significant group differences in network coherence (see previous section). We found that SI+ did not differ from the other two groups in between-network connectivity. In contrast, NSSI+ had significantly higher connectivity between the CEN networks and DMN-P compared to both NSSI- and CTL, as well as significantly higher connectivity between CEN-R and DMN-V compared to NSSI-(all *p*s<0.030). The NSSI-group did not differ from CTL in any CEN– DMN connectivity (all *p*s>0.119). See Figure 4 and “Between-Network Connectivity Results” in the Supplement for more details.

**Figure 4.** Between network connectivity associated with non-suicidal self-injury. NSSI+ exhibited significantly higher connectivity between the right **(A)** and left **(B)** CEN and DMN-P compared to NSSI- and CTL. NSSI+ exhibited significantly higher connected between the right CEN and DMN-V **(C)** compared to NSSI-but not CTL. CEN =central executive network; DMN-P=posterior default mode network; NSSI=non-suicidal self-injury; CTL=healthy controls \**p*<0.05, \*\**p*<0.01 (uncorrected)

### Correlations Between Network Coherence and Behavior

Among the depressed adolescents, higher SIQ and SITBI scores were significantly associated with lower coherence in the DMN-A, DMN-P, DMN-V, and SN-Ins (all *p*s<0.001). No other behavioral correlations we examined were significant. See “Correlations Between Network Coherence and Behavior” in the Supplement for more details.

## Discussion

The present investigation is the largest resting-state fMRI study examining the neural correlates of SI and NSSI in adolescents, the first to include clinical controls in the context of NSSI, and the first to identify subnetworks of the DMN and SN that are unique to NSSI. Importantly, although our sample of adolescents with SI and NSSI overlapped substantially, those with suicidal ideation (SI+) did not differ from those without suicidal ideation (SI-) in history or frequency of NSSI, medication status, or depression severity; similarly, adolescents with NSSI (NSSI+) did not differ from those without NSSI (NSSI-) in history of suicide attempt, history of SI, severity of SI, medication status, or depression severity. An additional strength of our study is our use of a multivariate data-driven approach to characterize intrinsic networks in order to resolve subnetworks of the DMN and SN, which are the two primary networks for which investigators have reported inconsistent patterns of resting-state functional alterations associated with SI and NSSI in adolescents (11). Finally, we complemented our within-network connectivity analyses with between-network connectivity analyses. We found that: 1) SI+ differed significantly from the other two groups in network coherence in the DMN-P; 2) NSSI+ differed significantly from the other two groups in network coherence in the DMN-A, DMN-V, and SN-Ins; 3) NSSI+ exhibited greater CEN connectivity with DMN-P compared to NSSI-. Our investigation identifies, for the first time, the within- and between-network patterns of connectivity associated with SI and NSSI in depressed adolescents; these findings offer insight concerning which intrinsic functional networks underlie SI and NSSI in depressed adolescents and inform future research that seeks to understand the etiology and treatment of these co-occurring behaviors.

We found that SI+ had lower DMN-V coherence than did both SI- and healthy controls, whereas both SI+ and SI-each had lower network coherence in DMN-P than did healthy controls (SI+ and SI-did not differ in DMN-P coherence). Similarly, we found that NSSI+ had lower DMN-V coherence than did both NSSI- and healthy controls, whereas both NSSI+ and NSSI-had lower network coherence in DMN-P than did healthy controls (NSSI+ and NSSI-did not differ in DMN-P coherence). NSSI+ also had lower CEN–DMN-V connectivity compared to NSSI-. Together, these results suggest that whereas lower DMN-V coherence is specifically related to thoughts and actions of harming one’s self, lower coherence of the DMN-P may be a more general marker of depression. Indeed, several studies with both depressed adolescents and adults have reported altered DMN-P connectivity compared to healthy controls (55, 56, 57). Other studies have also demonstrated that the DMN-P is involved in self-referential processing, particularly self-relevant affective information (34, 58).

In contrast, the DMN-V (which in our sample was centered in the middle temporal lobe, hippocampus, and parahippocampal region) has been shown in studies of healthy young adults to be a distinct subsystem of the DMN that is specifically involved in mnemonic scene construction and imagining one’s future (34, 59, 60). Interestingly, activation in the middle temporal lobe specifically during reflection of death- and life-related concepts has also been previously identified as one of the most highly discriminative brain patterns between young adults with suicidal ideation and healthy controls when using machine learning (61). While we were unable to explicitly relate DMN-V coherence with instructions to reflect on one’s future self, our findings are consistent with the formulation that suicidal ideation and non-suicidal self-injury are, in part, characterized by difficulties in imagining and planning for one’s future. Future work is needed to test whether psychosocial therapies that target representations of the self and that engage the DMN-V are effective in treating adolescents experiencing SI and NSSI (13, 31).

We also found that NSSI+ exhibited lower DMN-A and SN-Ins coherence compared to both NSSI- and healthy controls; SI+ and SI-adolescents did not differ significantly in coherence of these networks, suggesting that these patterns are specific to NSSI. Several studies of adults with Borderline Personality Disorder and NSSI have shown altered DMN and SN connectivity in relation to pain processing (62, 63; for reviews, see 12 and 64). It is notable that several fMRI studies have also identified altered connectivity of the DMN-P in patients with NSSI, but they did not include clinical controls; thus, it is possible that altered DMN-P connectivity is a depression-general marker, not a marker specific to NSSI. The DMN-A—and, specifically, the medial PFC, and its role in self-evaluative processes, particularly of the present self, may potentially represent an important neurobiological target for effective treatment of NSSI in adolescents. However, future research is needed to test these possibilities.

Our finding of NSSI+ exhibiting lower SN-Ins coherence relative to both NSSI- and healthy controls is highly consistent with previous studies of transdiagnostic samples of adolescents with NSSI that have identified lower insula-based resting-state functional connectivity (31). The SN-Ins network derived in our sample was centered primarily in the insula, striatum, and somatosensory regions. Both the insula and somatosensory cortex have been implicated in the processing of painful stimuli in studies of adults with BPD and/or NSSI (65, 66). Converging lines of evidence have pointed to the critical role of the insula, particularly in the posterior and mid-insular regions, in interoceptive awareness (67, 68), whereas the anterior insula (which in our sample was a core region of the SN-ACC) appears to be implicated more strongly in emotional awareness and in the integration of affect in decision-making, particularly with negatively valenced stimuli (69, 70). The insula is also strongly connected with sensorimotor and somatosensory regions; reduced connectivity among these regions may therefore reflect lower pain sensitivity and altered pain perception in adolescents with NSSI (12, 64). Together, our findings highlight an important direction for future research focused on understanding connectivity between the insula and somatosensory regions in adolescents with NSSI.

Contrary to our hypotheses, which were guided by previously published literature, we did not find group differences with respect to SI history in network coherence or between-network connectivity of the CEN or SN-ACC. Previous work from our group using similar analytical approaches (i.e., group ICA combined with dual regression) in an independent sample of depressed adolescents (24, 25) found that lower CEN, DMN-A, and SN-ACC coherence levels were associated with greater intensity of the most severe experience of SI in the lifetime (24), and that longitudinal changes in SN-ACC coherence were associated with changes in SI severity (25). Another study also reported that compared to healthy controls, young adults (who were primarily diagnosed with depression) with SI had lower dorsal ACC low frequency BOLD activity and relative connectivity (with dorsal versus ventral PCC; Chase et al., 2017). However, in our group’s previous investigations (24, 25), we had not recruited a psychiatrically healthy comparison group; moreover, all three of these studies did not include clinical controls. Hyperconnectivity between the CEN and DMN-P (the DMN most strongly associated with rumination) may reflect inadequate engagement of cognitive control due to the dominance of negative self-evaluative thinking (56, 57, 71). Given the extensive literature documenting differences in seed-based connectivity in regions of the CEN and SN-ACC in depressed adolescents and adults (55, 57) our results suggest that brain connectivity patterns associated with suicidal ideation may be specific to the DMN. Consistent with this formulation is the fact that we found that higher severity of suicidal ideation in the past month was significantly associated with lower coherence in all three DMNs.

While we did not find group differences with respect to NSSI history in network coherence of the CEN or SN-ACC, we did find that depressed adolescents with NSSI exhibited higher connectivity between the CEN and DMN-P compared to both those without history of NSSI and healthy controls. Previous resting-state fMRI studies have only compared transdiagnostic samples of adolescents with NSSI to healthy controls (or have conducted within-individual analyses in the context of a treatment study); in these studies, adolescents with NSSI were characterized by lower connectivity of the SN, as well as the DMN (31, 32, 33). Our study therefore clarifies the role of DMN connectivity in NSSI specifically by demonstrating that both within- and between-network connectivity of the DMN is compromised in adolescents with NSSI.

Because our primary aims were to examine differences in network coherence between SI+ and NSSI+ compared to both clinical and healthy controls, to ensure comparability across participants, and to increase generalizability of our findings while reliably estimating individual-level metrics of network coherence, we elected to conduct group-ICA followed by dual regression. For resting-state fMRI scans to have clinical utility, however, individual-level networks must ultimately be generated. While the consensus in the field is that longer scan times yield more reliable estimates of individual-level networks (72, 73), resting-state fMRI scans that are 5-7 minutes represent a pragmatic tradeoff between scan acquisition length and robustness of intrinsic functional networks, particularly in pediatric samples (74, 75). Nevertheless, previous studies have demonstrated that subsets of individuals may exhibit network components that are not found in group-levels maps (e.g., portions of medial PFC in the SN rather than the DMN; 76, 77). Thus, future work will benefit from examining individualized intrinsic networks derived from longer scans.

Future research is also needed to address the limitations of our current investigation. The SI and NSSI groups were highly overlapping in our sample of depressed adolescents; these rates, however, matched those typically seen in clinical populations (6, 7), thereby increasing the generalizability of our findings. Importantly, the depressed adolescents with and without history of SI did not differ significantly in history or frequency of NSSI. Similarly, the depressed adolescents with and without history of NSSI did not differ significantly in history of suicide attempt or in history or severity of SI. Thus, our study design was able to identify dissociable brain network patterns specific to SI+ and NSSI+ relative to SI- and NSSI-comparisons, respectively. Nevertheless, it will be important for future research to continue to work to dissociate the neural correlates of SI and NSSI by recruiting at least 4 independent groups presenting with all possible combinations of these behaviors to examine their shared and unique neurobiological substrates.

In addition, our investigation is limited by the exclusive focus on depressed adolescents. While studying a sample of adolescents with depressive disorders reduces clinical heterogeneity, the neural patterns identified in the current study may not be representative of the neural circuits underpinning SI and NSSI in adolescents more broadly; thus, it will be important in future studies to examine SI and NSSI transdiagnostically. Longitudinal studies with larger samples are also needed to clarify which brain markers are associated with past history, current indicators of SI and NSSI, and risk for the subsequent development of these behaviors. We will continue to follow this sample to determine whether network coherence (or between-network connectivity) predicts subsequent SI and NSSI. Another important limitation of our study is that we did not assess constructs related to self-referential thinking (e.g., imagining one’s future self, rumination) or interoceptive awareness and were thus unable to relate these constructs with their purported neural underpinnings in the intrinsic networks we assessed. Finally, the neurobiological interpretation of many network-based analyses, including ICA, remains unknown. Other network-based approaches, including graph theory, may be useful in providing complementary information about the organizational properties of specific brain regions in the context of large-scale networks; indeed, graph metrics have high test re-test reliability (78) and contain organizational properties (e.g., number and strength of connections among brain regions) that can augment our current understanding of network-based disruptions associated with SI and NSSI. However, parcellation decisions remain a challenge for the application of graph theory to neuroimaging data (79); ICA may represent a helpful first step for narrowing the parameter space for investigations computing graph metrics. Ultimately, targeted investigations that probe and characterize activation and connectivity of specific brain regions using multimodal approaches will be needed to enhance our understanding of network-based neural patterns that underlie SI and NSSI in adolescents.

In summary, we present new evidence that SI and NSSI are both characterized by lower connectivity in the ventral DMN, suggestive of difficulties in self-referential processing in the context of imagining and planning for the future, and that NSSI specifically is characterized by lower connectivity in the anterior DMN and in the insula and somatosensory network, suggestive of disruptions in self-evaluative processing, perception of sensory stimuli, and integration of interoceptive awareness and bodily signals. Our findings can guide future research seeking to understand the neurobiological underpinnings of SI and NSSI in adolescents.

## Acknowledgments

This research was supported by the National Institutes of Health (K01MH117442 to TCH, R37MH101495 to IHG), the Klingenstein Third Generation Foundation (Child and Adolescent Depression Fellow Award to TCH), the Stanford Maternal Child and Health Institute (Early Career Award and K Support Award to TCH), the Stanford Center for Cognitive and Neurobiological Imaging Center (Seed Grant to TCH), and the Ray and Dagmar Dolby Family Fund (to TCH). The funding agencies played no role in the design and conduct of the study; collection, management, analysis, and interpretation of the data; and preparation, review, or approval of the manuscript. MKS serves on the advisory board for Sunovion, is a consultant for Google X and Limbix, and receives royalties from the American Psychiatric Association Publishing. All other authors report no biomedical conflicts of interest. We thank Anna Cichocki, Miranda Edwards, Amar Ojha, Holly Pham, Michelle Sanabria, Lucinda Sisk, Jillian Segarra, Alexess Sosa, and Rachel Weisenburger for assistance with data collection and organization. Finally, we thank the participants and their families for their time and dedication in participating in this study.

## References

1. Curtin SC (2019): Death Rates Due to Suicide and Homicide Among Persons Aged 10–24: United States, 2000–2017. 8.

2. Nock MK, Green JG, Hwang I, McLaughlin KA, Sampson NA, Zaslavsky AM, Kessler RC (2013): Prevalence, Correlates, and Treatment of history Suicidal Behavior Among Adolescents: Results From the National Comorbidity Survey Replication Adolescent Supplement. JAMA Psychiatry 70: 300–310. 2.

3. Plener PL, Schumacher TS, Munz LM, Groschwitz RC (2015): The longitudinal course of non-suicidal self-injury and deliberate self-harm: a systematic review of the literature. Borderline Personal Disord Emot Dysregul 2: 2.

4. Franklin JC, Ribeiro JD, Fox KR, Bentley KH, Kleiman EM, Huang X, et al. (2017): Risk factors for suicidal thoughts and behaviors: A meta-analysis of 50 years of research. Psychological Bulletin 143: 187–232.

5. Gillies D, Christou MA, Dixon AC, Featherston OJ, Rapti I, Garcia-Anguita A, et al. (2018): Prevalence and Characteristics of Self-Harm in Adolescents: Meta-Analyses of Community-Based Studies 1990–2015. Journal of the American Academy of Child & Adolescent Psychiatry 57: 733–741.

6. Asarnow JR, Porta G, Spirito A, Emslie G, Clarke G, Wagner KD, et al. (2011): Suicide Attempts and Nonsuicidal Self-Injury in the Treatment of Resistant Depression in Adolescents: Findings from the TORDIA Study. Journal of the American Academy of Child & Adolescent Psychiatry 50: 772–781.

7. Glenn CR, Lanzillo EC, Esposito EC, Santee AC, Nock MK, Auerbach RP (2017): Examining the Course of Suicidal and Nonsuicidal Self-Injurious Thoughts and Behaviors in Outpatient and Inpatient Adolescents. J Abnorm Child Psychol 45: 971–983.

8. Hamza CA, Stewart SL, Willoughby T (2012): Examining the link between nonsuicidal self-injury and suicidal behavior: A review of the literature and an integrated model. Clinical Psychology Review 32: 482–495.

9. Liu RT (2017): Characterizing the course of non-suicidal self-injury: A cognitive neuroscience perspective. Neuroscience & Biobehavioral Reviews 80: 159–165.

10. Brown RC, Plener PL (2017): Non-suicidal Self-Injury in Adolescence. Curr Psychiatry Rep 19: 20.

11. Auerbach RP, Pagliaccio D, Allison GO, Alqueza KL, Alonso MF (2020) Neural Correlates Associated with Suicide and Non-Suicidal Self-Injury in Youth. Biological Psychiatry.

12. Westlund Schreiner M, Klimes-Dougan B, Begnel ED, Cullen KR (2015): Conceptualizing the neurobiology of non-suicidal self-injury from the perspective of the Research Domain Criteria Project. Neuroscience & Biobehavioral Reviews 57: 381–391.

13. Westlund Schreiner M, Klimes-Dougan B, Parenteau A, Hill D, Cullen KR (2019): A Framework for Identifying Neurobiologically Based Intervention Targets for NSSI. Curr Behav Neurosci Rep 6: 177–187.

14. Schmaal L, van Harmelen A-L, Chatzi V, Lippard ETC, Toenders YJ, Averill LA, et al. (2020): Imaging suicidal thoughts and behaviors: a comprehensive review of 2 decades of neuroimaging studies. Mol Psychiatry 25: 408–427.

15. Menon V (2011): Large-scale brain networks and psychopathology: a unifying triple network model. Trends in Cognitive Sciences 15: 483–506.

16. Whitfield-Gabrieli S, Ford JM (2012): Default Mode Network Activity and Connectivity in Psychopathology. Annual Review of Clinical Psychology 8: 49–76.

17. Berman MG, Peltier S, Nee DE, Kross E, Deldin PJ, Jonides J (2011): Depression, rumination and the default network. Soc Cogn Affect Neurosci 6: 548–555.

18. Hamilton JP, Farmer M, Fogelman P, Gotlib IH (2015): Depressive Rumination, the Default-Mode Network, and the Dark Matter of Clinical Neuroscience. Biological Psychiatry 78: 224–230.

19. Menon V, Uddin LQ (2010): Saliency, switching, attention and control: a network model of insula function. Brain Struct Funct 214: 655–667.

20. Seeley WW, Menon V, Schatzberg AF, Keller J, Glover GH, Kenna H, et al. (2007): Dissociable Intrinsic Connectivity Networks for Salience Processing and Executive Control. Journal of Neuroscience 27: 2349–2356.

21. Heuvel MP van den, Mandl RCW, Kahn RS, Pol HEH (2009): Functionally linked resting-state networks reflect the underlying structural connectivity architecture of the human brain. Human Brain Mapping 30: 3127–3141.

22. Thomas Yeo BT, Krienen FM, Sepulcre J, Sabuncu MR, Lashkari D, Hollinshead M, et al. (2011): The organization of the human cerebral cortex estimated by intrinsic functional connectivity. Journal of Neurophysiology 106: 1125–1165.

23. Schreiner MW, Klimes-Dougan B, Cullen KR (2019): Neural Correlates of Suicidality in Adolescents with Major Depression: Resting-State Functional Connectivity of the Precuneus and Posterior Cingulate Cortex. Suicide and Life-Threatening Behavior 49: 899–913.

24. Ordaz SJ, Goyer MS, Ho TC, Singh MK, Gotlib IH (2018): Network basis of suicidal ideation in depressed adolescents. J Affect Disord 226: 92–99.

25. Schwartz J, Ordaz SJ, Ho TC, Gotlib IH (2019): Longitudinal decreases in suicidal ideation are associated with increases in salience network coherence in depressed adolescents. Journal of Affective Disorders 245: 545–552.

26. Chase HW, Segreti AM, Keller TA, Cherkassky VL, Just MA, Pan LA, Brent DA (2017): Alterations of functional connectivity and intrinsic activity within the cingulate cortex of suicidal ideators. Journal of Affective Disorders 212: 78–85.

27. Johnston JAY, Wang F, Liu J, Blond BN, Wallace A, Liu J, et al. (2017): Multimodal Neuroimaging of Frontolimbic Structure and Function Associated With Suicide Attempts in Adolescents and Young Adults With Bipolar Disorder. AJP 174: 667–675.

28. Zhang S, Chen J, Kuang L, Cao J, Zhang H, Ai M, et al. (2016): Association between abnormal default mode network activity and suicidality in depressed adolescents. BMC Psychiatry 16: 337.

29. Cao J, Ai M, Chen X, Chen J, Wang W, Kuang L (2020): Altered resting-state functional network connectivity is associated with suicide attempt in young depressed patients. Psychiatry Research 285: 112713.

30. Qiu H, Cao B, Cao J, Li X, Chen J, Wang W, et al. (2020): Resting-state functional connectivity of the anterior cingulate cortex in young adults depressed patients with and without suicidal behavior. Behavioural Brain Research 384: 112544.

31. Santamarina-Perez P, Romero S, Mendez I, Leslie SM, Packer MM, Sugranyes G, et al. (2019): Fronto-Limbic Connectivity as a Predictor of Improvement in Nonsuicidal Self-Injury in Adolescents Following Psychotherapy. Journal of Child and Adolescent Psychopharmacology 29: 456–465.

32. Westlund Schreiner M, Klimes-Dougan B, Mueller BA, Eberly LE, Reigstad KM, Carstedt PA, et al. (2017): Multi-modal neuroimaging of adolescents with non-suicidal self-injury: Amygdala functional connectivity. Journal of Affective Disorders 221: 47–55.

33. Cullen KR, Schreiner MW, Klimes-Dougan B, Eberly LE, LaRiviere LL, Lim KO, et al. (2020): Neural correlates of clinical improvement in response to N-acetylcysteine in adolescents with non-suicidal self-injury. Progress in Neuro-Psychopharmacology and Biological Psychiatry 99: 109778.

34. Andrews-Hanna JR, Reidler JS, Sepulcre J, Poulin R, Buckner RL (2010): Functional-Anatomic Fractionation of the Brain’s Default Network. Neuron 65: 550–562.

35. Chong JSX, Ng GJP, Lee SC, Zhou J (2017): Salience network connectivity in the insula is associated with individual differences in interoceptive accuracy. Brain Struct Funct 222: 1635–1644.

36. Dixon ML, Vega ADL, Mills C, Andrews-Hanna J, Spreng RN, Cole MW, Christoff K (2018): Heterogeneity within the frontoparietal control network and its relationship to the default and dorsal attention networks. PNAS 115: E1598–E1607.

37. Greicius MD, Supekar K, Menon V, Dougherty RF (2009): Resting-state functional connectivity reflects structural connectivity in the default mode network. Cereb Cortex 19: 72–78.

38. Zuo X-N, Kelly C, Adelstein JS, Klein DF, Castellanos FX, Milham MP (2010): Reliable intrinsic connectivity networks: Test–retest evaluation using ICA and dual regression approach. NeuroImage 49: 2163–2177.

39. Smith DV, Utevsky AV, Bland AR, Clement N, Clithero JA, Harsch AEW, et al. (2014): Characterizing individual differences in functional connectivity using dual-regression and seed-based approaches. NeuroImage 95: 1–12.

40. Kaufman J, Birmaher B, Brent D, Rao U, Flynn C, Moreci P, et al. (1997): Schedule for Affective Disorders and Schizophrenia for School-Age Children-Present and Lifetime Version (K-SADS-PL): Initial Reliability and Validity Data. Journal of the American Academy of Child & Adolescent Psychiatry 36: 980–988.

41. Kaufman J, Birmaher B, Brent DA, Ryan, ND, & Rao U (2000). K-SADS-PL. Journal of the American Academy of Child & Adolescent Psychiatry, 39(10): 1208. https://doi.org/10.1097/00004583-200010000-00002

42. Poznanski EO, & Mokros HB (1996) Children’s Depression Rating Scale, Revised (CDRS-R). Los Angeles: Western Psychological Services.

43. Maxwell ME (1992). Family Interview for Genetic Studies (FIGS): a manual for FIGS. Bethesda, MD: Clinical Neurogenetics Branch, Intramural Research Program, National Institute of Mental Health.

44. Posner K, Brown GK, Stanley B, Brent DA, Yershova KV, Oquendo MA, et al. (2011): The Columbia-Suicide Severity Rating Scale: initial validity and internal consistency findings from three multisite studies with adolescents and adults. Am J Psychiatry 168: 1266–1277.

45. Nock MK, Holmberg EB, Photos VI, Michel BD (2007): Self-Injurious Thoughts and Behaviors Interview: Development, reliability, and validity in an adolescent sample. Psychological Assessment 19: 309–317.

46. Reynolds WM (1987): Suicidal ideation questionnaire (SIQ). Odessa, FL: Psychological Assessment Resources.

47. Schwartz J, Ordaz SJ, Kircanski K, Ho TC, Davis EG, Camacho MC, Gotlib IH (2019): Resting-state functional connectivity and inflexibility of daily emotions in major depression. Journal of Affective Disorders 249: 26–34.

48. Ordaz SJ, LeMoult J, Colich NL, Prasad G, Pollak M, Popolizio M, et al. (2017): Ruminative brooding is associated with salience network coherence in early pubertal youth. Soc Cogn Affect Neurosci 12: 298–310.

49. Fischl B (2012): FreeSurfer. Neuroimage 62: 774–781.

50. Smith SM, Jenkinson M, Woolrich MW, Beckmann CF, Behrens TEJ, Johansen-Berg H, et al. (2004): Advances in functional and structural MR image analysis and implementation as FSL. Neuroimage 23 Suppl 1: S208–219.

51. Cox RW (1996): AFNI: software for analysis and visualization of functional magnetic resonance neuroimages. Comput Biomed Res 29: 162–173.

52. Beckmann, C. F., Mackay, C. E., Filippini, N., & Smith, S. M. (2009). Group comparison of resting-state FMRI data using multi-subject ICA and dual regression. Neuroimage, 47(Suppl 1), S148.

53. Kiviniemi V, Kantola J-H, Jauhiainen J, Hyvärinen A, Tervonen O (2003): Independent component analysis of nondeterministic fMRI signal sources. NeuroImage 19: 253–260.

54. Bunn A, Korpela M (n.d.): An Introduction to dplR. 16.

55. Kerestes R, Davey CG, Stephanou K, Whittle S, Harrison BJ (2014): Functional brain imaging studies of youth depression: A systematic review. NeuroImage: Clinical 4: 209–231.

56. Ho TC, Connolly CG, Henje Blom E, LeWinn KZ, Strigo IA, Paulus MP, et al. (2015): Emotion-Dependent Functional Connectivity of the Default Mode Network in Adolescent Depression. Biological Psychiatry 78: 635–646.

57. Kaiser RH, Andrews-Hanna JR, Wager TD, Pizzagalli DA (2015): Large-Scale Network Dysfunction in Major Depressive Disorder: A Meta-analysis of Resting-State Functional Connectivity. JAMA Psychiatry 72: 603–611.

58. Davey CG, Pujol J, Harrison BJ (2016): Mapping the self in the brain’s default mode network. NeuroImage 132: 390–397.

59. Bellana B, Liu Z-X, Diamond NB, Grady CL, Moscovitch M (2017): Similarities and differences in the default mode network across rest, retrieval, and future imagining. Human Brain Mapping 38: 1155–1171.

60. Schacter DL, Addis DR, Hassabis D, Martin VC, Spreng RN, Szpunar KK (2012): The Future of Memory: Remembering, Imagining, and the Brain. Neuron 76: 677–694.

61. Just MA, Pan L, Cherkassky VL, McMakin D, Cha C, Nock MK, Brent D (2017): Machine learning of neural representations of suicide and emotion concepts identifies suicidal youth. Nat Hum Behav 1: 911–919.

62. Kluetsch RC, Schmahl C, Niedtfeld I, Densmore M, Calhoun VD, Daniels J, et al. (2012): Alterations in default mode network connectivity during pain processing in borderline personality disorder. Arch Gen Psychiatry 69: 993–1002.

63. Osuch E, Ford K, Wrath A, Bartha R, Neufeld R (2014): Functional MRI of pain application in youth who engaged in repetitive non-suicidal self-injury vs. psychiatric controls. Psychiatry Research: Neuroimaging 223: 104–112.

64. Ballard E, Bosk A, Pao M (2010): Invited Commentary: Understanding Brain Mechanisms of Pain Processing in Adolescents’ Non-Suicidal Self-Injury. J Youth Adolescence 39: 327–334.

65. Bonenberger M, Plener PL, Groschwitz RC, Grön G, Abler B (2015): Differential neural processing of unpleasant haptic sensations in somatic and affective partitions of the insula in non-suicidal self-injury (NSSI). Psychiatry Research: Neuroimaging 234: 298–304.

66. Niedtfeld I, Schmitt R, Winter D, Bohus M, Schmahl C, Herpertz SC (2017): Pain-mediated affect regulation is reduced after dialectical behavior therapy in borderline personality disorder: a longitudinal fMRI study. Soc Cogn Affect Neurosci 12: 739–747.

67. Khalsa SS, Rudrauf D, Feinstein JS, Tranel D (2009): The pathways of interoceptive awareness [no. 12]. Nature Neuroscience 12: 1494–1496.

68. Kurth F, Eickhoff SB, Schleicher A, Hoemke L, Zilles K, Amunts K (2010): Cytoarchitecture and probabilistic maps of the human posterior insular cortex. Cereb Cortex 20: 1448–1461.

69. Caria A, Sitaram R, Veit R, Begliomini C, Birbaumer N (2010): Volitional Control of Anterior Insula Activity Modulates the Response to Aversive Stimuli. A Real-Time Functional Magnetic Resonance Imaging Study. Biological Psychiatry 68: 425–432.

70. Singer T, Critchley HD, Preuschoff K (2009): A common role of insula in feelings, empathy and uncertainty. Trends in Cognitive Sciences 13: 334–340.

71. Jacobs RH, Jenkins LM, Gabriel LB, Barba A, Ryan KA, Weisenbach SL, et al. (2014): Increased Coupling of Intrinsic Networks in Remitted Depressed Youth Predicts Rumination and Cognitive Control. PLoS One 9. https://doi.org/10.1371/journal.pone.0104366

72. Birn RM, Molloy EK, Patriat R, Parker T, Meier TB, Kirk GR, et al. (2013): The effect of scan length on the reliability of resting-state fMRI connectivity estimates. NeuroImage 83: 550–558.

73. Noble S, Spann MN, Tokoglu F, Shen X, Constable RT, Scheinost D (2017): Influences on the Test–Retest Reliability of Functional Connectivity MRI and its Relationship with Behavioral Utility. Cereb Cortex 27: 5415–5429.

74. Van Dijk KRA, Hedden T, Venkataraman A, Evans KC, Lazar SW, Buckner RL (2009): Intrinsic Functional Connectivity As a Tool For Human Connectomics: Theory, Properties, and Optimization. Journal of Neurophysiology 103: 297–321.

75. Whitlow CT, Casanova R, Maldjian JA (2011): Effect of Resting-State Functional MR Imaging Duration on Stability of Graph Theory Metrics of Brain Network Connectivity. Radiology 259: 516–524.

76. Gordon EM, Laumann TO, Gilmore AW, Newbold DJ, Greene DJ, Berg JJ, et al. (2017): Precision Functional Mapping of Individual Human Brains. Neuron 95: 791-807.e7.

77. Tavor I, Jones OP, Mars RB, Smith SM, Behrens TE, Jbabdi S (2016): Task-free MRI predicts individual differences in brain activity during task performance. Science 352: 216–220.

78. Yuan JP, Henje Blom E, Flynn T, Chen Y, Ho TC, Connolly CG, et al. (2019): Test–Retest Reliability of Graph Theoretic Metrics in Adolescent Brains. Brain Connectivity 9: 144–154.

79. Ho TC, Dennis EL, Thompson PM, Gotlib IH (2018): Network-based approaches to examining stress in the adolescent brain. Neurobiology of Stress 8: 147–157.

