## Supplement for "Default Mode and Salience Network Alterations in Suicidal and Non-Suicidal Self-Injurious Thoughts and Behaviors in Adolescents with Depression"

**Supplemental Information to “Default Mode and Salience Network Alterations in Suicidal Ideation and Non-Suicidal Self-Injury in Adolescents with Depression”**

*Clinical Assessments*

We interviewed participants at an initial behavioral session to assess study eligibility using the Kiddie Schedule for Affective Disorders and Schizophrenia–Present and Lifetime (K-SADS-PL; 1, 2), Children’s Depressive Rating Scale–Revised (CDRS-R; 3), and Family Interview for Genetics Studies (4). For the present study, we administered Screening Interviews and Supplements from the K-SADS-PL for the following modules to determine study inclusion: Major Depressive Disorder, Mania, Psychosis, Cigarette/Tobacco Use, Alcohol Abuse, and Substance Use Disorder (1,2,5). CDRS-R scores were also used for determining study inclusion; both child- and parent-report were integrated and reviewed by a clinically trained research team and integrated to generate a single summary score. Standardized *t*-scores ≥ 55 indicate that the presence of a depressive disorder is likely to be confirmed in a comprehensive diagnostic evaluation and, thus, was used as a cut-off for study eligibility for those who did not meet full criteria for MDD or Dysthymia in the K-SADS-PL screening, provided that the participant also endorsed at least 2 symptoms of MDD or Dysthymia in the K-SADS-PL screening.

*Inclusion/Exclusion Criteria*

Inclusion criteria for potentially non-depressed adolescents included no current or lifetime diagnosis of any DSM-IV disorder or first-degree relative with history of suicide or known suicide attempt, or a suspected diagnosis of depression, bipolar disorder, or schizophrenia using the Family Interview for Genetics Study (4). Exclusion criteria were premenarchal status (for females), history of concussion within the past 6 weeks or history of any concussion with loss of consciousness, contraindications to MRI scanning (e.g. braces, metal implants, or claustrophobia), and any serious neurological or intellectual disorders that could interfere with the participant’s ability to complete study components; for potentially depressed adolescents, exclusion criteria also included meeting for lifetime or current criteria for Mania, Psychosis, or Alcohol Dependence (based on DSM-IV) or Moderate Substance Use Disorder with substance-specific threshold for withdrawal (based on DSM-V). One CTL participant met for a past episode of clinically significant SI but did not meet criteria for any DSM-IV diagnosis. We therefore conservatively included this participant in our analyses; as expected, removing them did not change the significance of any of our findings.

*MRI Scanning Acquisition*

All MRI scans were acquired at the Stanford Center for Cognitive and Neurobiological Imaging (CNI) with a 3T GE Discovery MR750 (General Electric Healthcare, Milwaukee, WI, USA) and Nova 32-channel head coil (Nova Medical, Wilmington, MA, USA). Participants completed a T2*weighted resting-state scan obtained using an echo planar image (EPI) sequence: 240 volumes, TR/TE=2000/30 ms; flip angle=77°; 42 oblique slices; FOV=240 mm; matrix=80 mm x 80 mm, voxel resolution=3x3x3 mm^3^; total scan time=8 min. During this scan, participants were instructed to close their eyes but remain awake. Participants also completed a T1-weighted anatomical scan obtained using a spoiled gradient (SPGR) sequence: TR/TE/TI=8.2/3.2/600 ms; flip angle=12°; 156 axial slices; FOV=256 mm; matrix=256 mm x 256 mm, voxel resolution=1x1x1 mm^3^; total scan time=3 min 40 sec. Physiological signal (i.e., respiration and heart rate) was collected via a photoplethysmograph attached to the toe and used for preprocessing the fMRI data (see next section).

*Resting-State fMRI Preprocessing*

All data were visually inspected prior to preprocessing to assess for magnetic field inhomogeneities or disruptions in the scanner signal. The SPGR images were skull-stripped and white matter (WM) and cerebrospinal fluid (CSF) were parcellated and eroded using Freesurfer’*s recon-all* routine (v. 6.0) before being registered to the MNI 2 mm^3^ standardized atlas using linear transformations in FSL (*FLIRT*). The appropriate WM and CSF erosion mask (1 voxel, 2 voxels, or 3 voxels) was selected for each participant based on visual inspection. The first six volumes of the functional time series were removed, leaving 234 volumes for subsequent preprocessing. Functional data were de-spiked (*3dDespike*) and physiological correction was applied (*RETROICOR*; 6, 7) before being slice-time corrected and spatially normalized to the MNI 2 mm^3^ standardized atlas (*3dAllineate*) and spatially smoothed using a 5 mm full width at half maximum Gaussian kernel (*3dBlurInMask*). Preprocessed data were then entered into a voxel-wise generalized least squares regression (*3dREMLfit*) in order to regress out spurious activation related to physiology and motion by including the mean WM timeseries, mean CSF timeseries and 24 motion parameters (3 translational, 3 rotational at each time point and the prior time point, and each of these 12 motion parameters squared) as covariates of non-interest. The resulting residuals were bandpass filtered (0.009 < *f* < 0.08 Hz; *3dBandpass*), de-meaned (*3dcalc*) and used for subsequent analyses (see “Defining Functional Networks Using Group Independent Component Analysis” in the Methods section of the main manuscript for more details). For each subject, we computed mean relative displacement (MRD, also known as framewise displacement) and excluded any participant with excessive movement, defined as having more than 20 volumes with a MRD > 0.25 mm or a mean MRD > 0.2 mm from subsequent analyses (8, 9). No participants moved excessively based on these thresholds; however, 7 participants (2 CTL) did not complete the full scan due to discomfort and 2 participants (1 CTL) did not have physiological signal properly recorded. Participants who were included in our final analyses did not differ from these 9 participants on any demographic characteristics (all *p*s>0.259).

*Between-Network Connectivity Results*

To complement our primary investigation examining network coherence (i.e., within-network connectivity), we tested whether groups differed in between-network connectivity in the CEN and SN-ACC, which were the networks where we did not find significant group differences in network coherence. SI+ did not differ from the other two groups in CEN or SN-ACC with one another or with the three DMN networks. In contrast, NSSI+ had significantly higher connectivity between CEN-L and DMN-P compared to both NSSI- (*B*=0.147±0.066, *t*(63)=2.217, *p*=0.030) and CTL (*B*=0.166±0.073, *t*(63)=2.261, *p*=0.027); NSSI+ also exhibited significantly higher connectivity between CEN-R and DMN-P compared to both NSSI- (*B*=0.213±0.071, *t*(63)=3.006, *p*=0.004) and CTL (*B*=0.177±0.079, *t*(63)=2.246, *p*=0.028). The NSSI- group, however, did not differ from CTL in connectivity between CEN and DMN-P (all *p*s>0.669). Finally, the NSSI+ group also exhibited significantly higher CEN-R connectivity with DMN-V compared to NSSI- (*B*=0.256±0.075, *t*(63)=3.410, *p*=0.001) but did not differ from CTL (*p*=0.119). See Figure 4 in the main text for more details.

*Correlations Between Network Coherence and Behavior*

Given the skewed distributions of 0the SIQ scores and NSSI frequency in the past month (see Figure S1), we conducted Poisson log-linear regressions to estimate the associations between coherence of DMN-A, DMN-V, DMN-P, and SN-Ins with SIQ scores and NSSI frequency in the past month. All statistical models included age, sex, motion, and medication usage as covariates. We found that higher SIQ scores were significantly associated with lower coherence in the DMN-A (*B*=-0.216±0.055, *z*(43)=-3.941, *p*=0.00008), DMN-P (*B*=-0.159±0.039, *z*(43)=-4.107, *p*=0.00004), DMN-V (*B*=-0.468±0.046, *z*(43)=-10.216, *p*=2x10^-16^), and SN-Ins (*B*=-0.295±0.043, *z*(43)=-6.808, *p*=9x10^-12^). See Figure S2 for more detail. Importantly, including current depression severity, as measured by CDRS-R *t*-scores, as a covariate in these models, did not alter the significance of our findings in the DMN-A, DMN-V, and SN-Ins (all *p*s<0.003); however, the association between SIQ scores and DMN-P did fall in significance after covarying for CDRS-R *t*-scores (*p*=0.060). Similarly, higher NSSI frequency in the past month was significantly associated with lower network coherence in all four of these networks (all *p*s<0.001); the significance of these associations did not change even when including current depression severity as a covariate (all *ps<*0.013).

**Table S1**. Full model results with covariates for each model with a significant effect of group (SI+, SI-, CTL) on network coherence. Sex and current medication were coded as binary factors. All covariates were mean-centered. See Table 2 in the main text for a summary of the group effect and linear contrast test results. All *p*-values reported are uncorrected. **p*<0.05, ***p*<0.01.

| **Model** | *Statistic* | *p* | *B*±SE | *Δ R^2^* | *Adj. R^2^* |
| --- | --- | --- | --- | --- | --- |
| *DMN-V Coherence* |  |  |  |  |  |
| Overall Model | *F*(6,63)=3.098 | 0.010* |  |  | 0.154 |
| Age | *t*(63)=0.615 | 0.541 | 0.041±0.067 | 0.006 |  |
| Sex | *t*(63)=-2.933 | 0.005** | -0.485±0.165 | 0.120 |  |
| Motion | *t*(63)=-1.958 | 0.055 | -8.216±4.196 | 0.057 |  |
| Current Meds | *t*(63)=-0.493 | 0.624 | -0.094±0.190 | 0.004 |  |
| *DMN-A Coherence* |  |  |  |  |  |
| Overall Model | *F*(6,63)=3.430 | 0.005** |  |  | 0.175 |
| Age | *t*(63)=1.328 | 0.189 | 0.077±0.058 | 0.027 |  |
| Sex | *t*(63)=-2.578 | 0.012* | -0.370±0.144 | 0.095 |  |
| Motion | *t*(63)=-1.342 | 0.184 | -4.893±3.647 | 0.028 |  |
| Current Meds | *t*(63)=0.986 | 0.328 | 0.163±0.165 | 0.015 |  |
| *DMN-P Coherence* |  |  |  |  |  |
| Overall Model | *F*(6,63)=2.723 | 0.020* |  |  | 0.130 |
| Age | *t*(63)=-1.365 | 0.177 | -0.112±0.082 | 0.029 |  |
| Sex | *t*(63)=-1.711 | 0.092 | -0.346±0.202 | 0.044 |  |
| Motion | *t*(63)=-2.018 | 0.048* | -10.364±5.137 | 0.061 |  |
| Current Meds | *t*(63)=-2.726 | 0.008** | -0.635±0.233 | 0.106 |  |
| *SN-Ins Coherence* |  |  |  |  |  |
| Overall Model | *F*(6,63)=2.589 | 0.026* |  |  | 0.121 |
| Age | *t*(63)=-1.069 | 0.289 | -0.089±0.083 | 0.018 |  |
| Sex | *t*(63)=-0.693 | 0.491 | -0.142±0.205 | 0.008 |  |
| Motion | *t*(63)=1.371 | 0.175 | 7.148±5.215 | 0.029 |  |
| Current Meds | *t*(63)=-0.890 | 0.377 | -0.211±0.237 | 0.012 |  |

**Table S2.** Full model results with covariates for each model with a significant effect of group (NSSI+, NSSI-, CTL) on network coherence. Sex and current medication were coded as binary factors. All covariates were mean-centered. See Table 3 in the main text for a summary of the group effect and linear contrast test results. All *p*-values reported are uncorrected. **p*<0.05, ***p*<0.01, ****p*<0.001.

| **Model** | *Statistic* | *p* | *B*±SE | *Δ R^2^* | *Adj. R^2^* |
| --- | --- | --- | --- | --- | --- |
| DMN-A Coherence |  |  |  |  |  |
| Overall Model | *F*(6,63)=4.682 | 0.0005*** |  |  | 0.243 |
| Age | *t*(63)=1.376 | 0.174 | 0.077±0.056 | 0.029 |  |
| Sex | *t*(63)=-2.766 | 0.007** | -0.381±0.138 | 0.108 |  |
| Motion | *t*(63)=-1.201 | 0.234 | -4.144±3.451 | 0.022 |  |
| Current Meds | *t*(63)=0.772 | 0.443 | 0.124±0.160 | 0.009 |  |
| DMN-V Coherence |  |  |  |  |  |
| Overall Model | *F*(6,63)=3.341 | 0.006** |  |  | 0.169 |
| Age | *t*(63)=0.610 | 0.544 | 0.041±0.067 | 0.006 |  |
| Sex | *t*(63)=-2.858 | 0.006** | -0.469±0.164 | 0.115 |  |
| Motion | *t*(63)=-1.507 | 0.137 | -6.191±4.108 | 0.035 |  |
| Current Meds | *t*(63)=-0.463 | 0.645 | -0.088±0.191 | 0.003 |  |
| SN-Ins Coherence |  |  |  |  |  |
| Overall Model | *F*(6,63)=4.356 | 0.001** |  |  | 0.226 |
| Age | *t*(63)=-1.020 | 0.311 | -0.080±0.078 | 0.016 |  |
| Sex | *t*(63)=-0.757 | 0.452 | -0.146±0.193 | 0.009 |  |
| Motion | *t*(63)=1.833 | 0.071 | 8.866±4.836 | 0.050 |  |
| Current Meds | *t*(63)=-0.925 | 0.359 | -0.208±0.224 | 0.013 |  |
| DMN-P Coherence |  |  |  |  |  |
| Overall Model | *F*(6,63)=3.439 | 0.005** |  |  | 0.175 |
| Age | *t*(63)=-1.439 | 0.155 | -0.115±0.080 | 0.032 |  |
| Sex | *t*(63)=-1.900 | 0.062 | -0.375±0.198 | 0.054 |  |
| Motion | *t*(63)=-2.070 | 0.043* | -10.230±4.942 | 0.064 |  |
| Current Meds | *t*(63)=-3.038 | 0.003** | -0.697±0.229 | 0.128 |  |

**Figure S1. Histograms of SIQ scores and frequency of NSSI (thoughts and actions) in the past month**. SIQ scores and past month NSSI frequency are from all depressed participants (*n*=49). SI=suicidal ideation; NSSI=non-suicidal self-injury; SIQ=Suicidal Ideation Questionnaire-Junior High; SITBI=Self-Injurious Thoughts and Behaviors Interview.

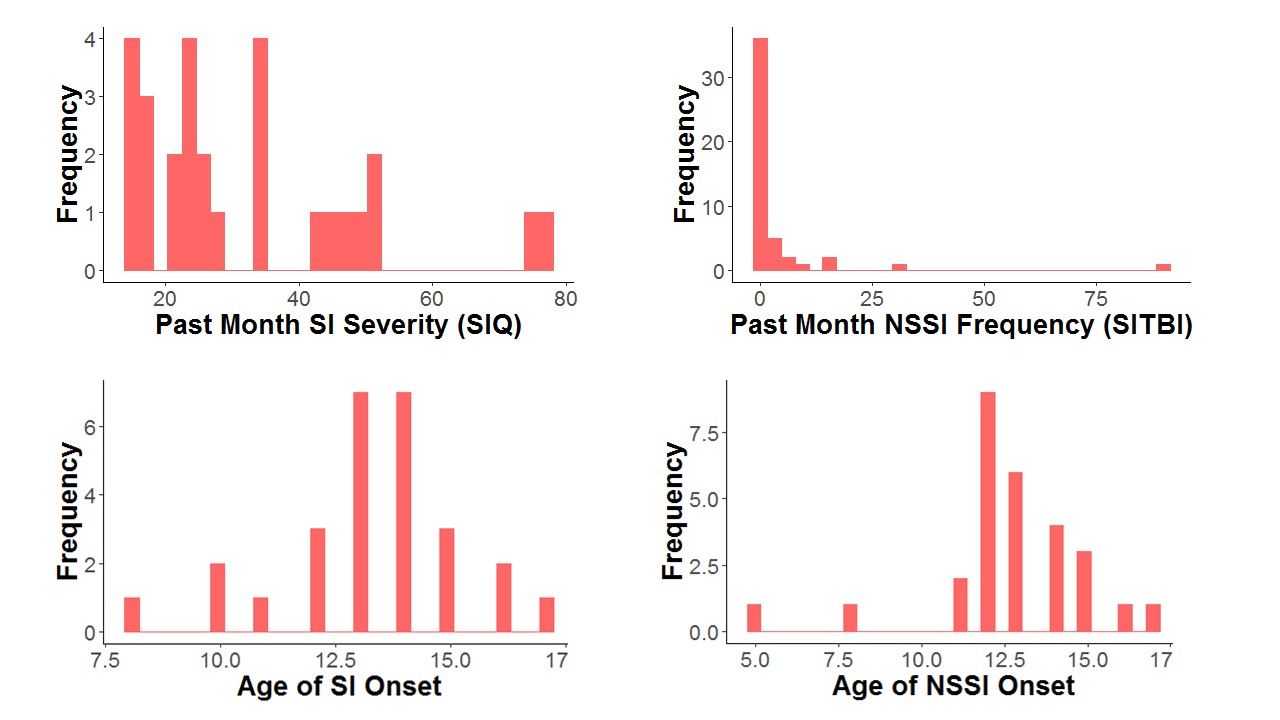

**Figure S2. Scatterplots of network coherence with SIQ scores.** Poisson log-linear regression models were used to examine associations between network coherence and past month suicidal ideation severity, as measured by SIQ scores. Lower coherence in the DMN-A, DMN-P, DMN-V, and SN-Ins were significantly associated with higher SIQ scores. DMN-A=anterior Default Mode Network; DMN-P=posterior Default Mode Network; DMN-V=ventral Default Mode Network; SN-Ins=insula-based Salience Network; SIQ=Suicidal Ideation Questionnaire-Junior High.

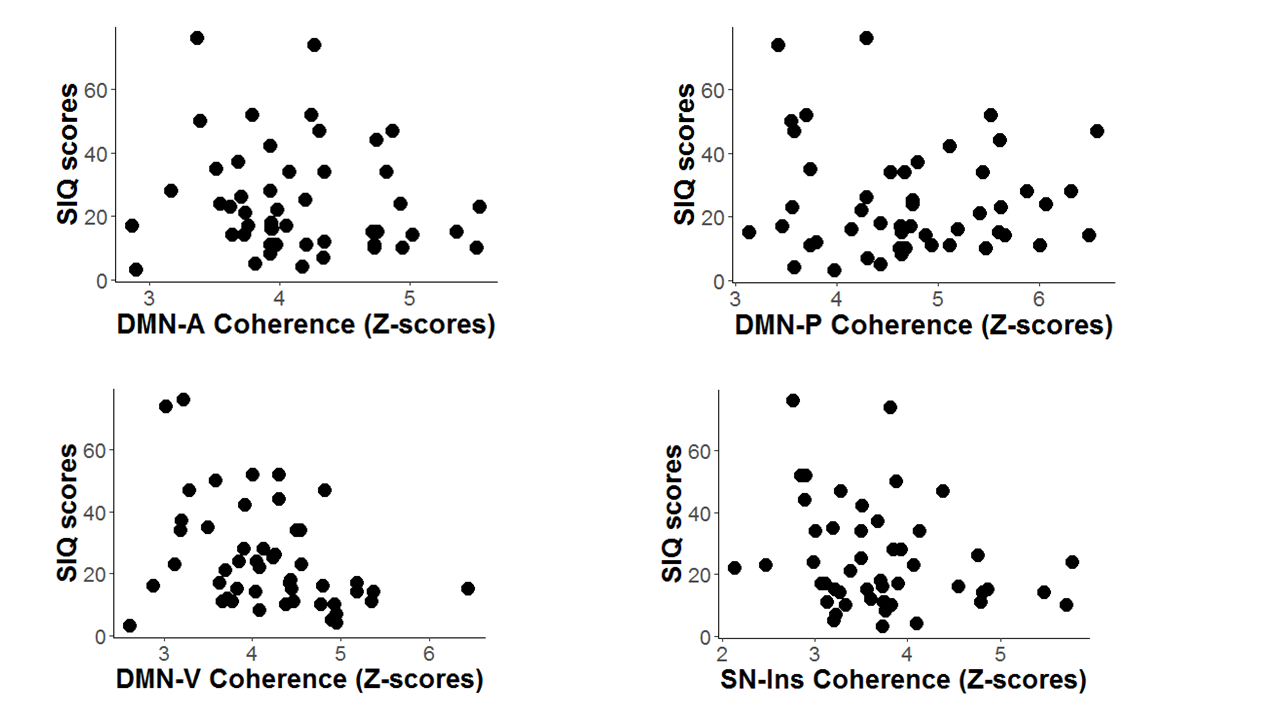

**Supplemental References**

1. Kaufman J, Birmaher B, Brent DA, Rao U, Flynn C, Moreci P, *et al.* (1997): Schedule for Affective Disorders and Schizophrenia for School-Age Children-Present and Lifetime Version (K-SADS-PL): Initial Reliability and Validity Data. *Journal of the American Academy of Child & Adolescent Psychiatry*, 36: 980–988.
2. Kaufman J, Birmaher B, Brent DA, Ryan, ND, & Rao U (2000). K-SADS-PL. *Journal of the American Academy of Child & Adolescent Psychiatry,* 39(10): 1208. [https://doi.org/10.1097/00004583-200010000-00002](https://psycnet.apa.org/doi/10.1097/00004583-200010000-00002)
3. Poznanski EO, & Mokros HB (1996) *Children's Depression Rating Scale, Revised (CDRS-R)*. Los Angeles: Western Psychological Services.
4. Maxwell ME (1992). Family Interview for Genetic Studies (FIGS): a manual for FIGS. *Bethesda, MD: Clinical Neurogenetics Branch, Intramural Research Program, National Institute of Mental Health*.
5. Birmaher B, Axelson D, Perepletchikova F, Brent DA, & Ryan N (2016) K-SADS-PL DSM-5. *Pittsburgh: Western Psychiatric Institute and Clinic*.
6. Chang C, Cunningham JP, & Glover GH (2009) Influence of heart rate on the BOLD signal: the cardiac response function, *Neuroimage*, 44(3):857–869.
7. Jo HJ, Saad ZS, Simmons WK, Milbury LA & Cox RW (2010) Mapping sources of correlation in resting state FMRI, with artifact detection and removal, *Neuroimage*, 52(2):571–582.
8. Satterthwaite TD, Elliott MA, Gerraty RT, Ruparel K, Loughead J, Calkins M, *et al.* (2013) An improved framework for confound regression and filtering for control of motion artifact in the preprocessing of resting-state functional connectivity data, *Neuroimage*, 64:240–256.
9. Power JD, Barnes KA, Snyder AZ, Schlaggar BL, & Petersen SE (2013) Steps toward optimizing motion artifact removal in functional connectivity MRI; a reply to Carp, *Neuroimage*, 76:439–441.
